# Bayesian multi-level calibration of a process-based maize phenology model

**DOI:** 10.1101/2022.06.03.494418

**Authors:** Michelle Viswanathan, Andreas Scheidegger, Thilo Streck, Sebastian Gayler, Tobias KD Weber

**Affiliations:** Institute of Soil Science and Land Evaluation, Biogeophysics, University of Hohenheim, Germany; Eawag: Swiss Federal Institute of Aquatic Science and Technology, Dubendorf, Switzerland

**Keywords:** phenology, silage maize, hierarchical Bayes, multi-level models, process-based model

## Abstract

Plant phenology models are important components in process-based crop models, which are used to assess the impact of climate change on food production. For reliable model predictions, parameters in phenology models have to be accurately known. They are usually estimated by calibrating the model to observations. However, at regional scales in which different cultivars of a crop species may be grown, not accounting for inherent differences in phenological development between cultivars in the model and the presence of model deficits lead to inaccurate parameter estimates. To account for inherent differences between cultivars and to identify model deficits, we used a Bayesian multi-level approach to calibrate a phenology model (SPASS) to observations of silage maize grown across Germany between 2009 and 2017. We evaluated four multi-level models of increasing complexity, where we accounted for different combinations of ecological, weather, and year effects, as well as the hierarchical classification of cultivars nested within ripening groups of the maize species. We compared the calibration quality from this approach to the commonly used pooled approach in which none of these factors are considered. Our approach proved successful in improving calibration quality by incorporating the hierarchical classification of cultivars within ripening groups of maize. Our findings have implications for regional model calibration and data-gathering studies, since it emphasizes that ripening group and cultivar information is essential. Furthermore, we found that if this information is not available, at least weather, ecological regions and year effects should be taken into account. Our results can facilitate model improvement studies since we identified possible model limitations related to temperature effects in the reproductive (post-flowering) phase and to soil-moisture. We demonstrate that Bayesian multi-level calibration of a phenology model facilitates the incorporation of hierarchical dependencies and the identification of model limitations. Our approach can be extended to full crop models at different spatial scales.

## 1 Introduction

Plant phenology plays an important role when evaluating the impact of climate change and assessing food security [1, 2, 3, 4, 5]. It is controlled by environmental variables and determines the timing of plant organ development and the distribution of assimilates to different parts of the plant. Thus, predictions of phenological development are essential for evaluating crop growth and yield, and may also support field management decisions such as the timing of fertilizer application [6]. These phenology predictions are made possible by using numerical models.

Phenology models are in turn important components of crop models, which are used for simulating crop growth and development, and yield. Besides data-driven statistical models, it is process-based models which enable a thorough understanding of the underlying processes for evaluating potential policy interventions and adaption to climate change [7]. In these process models, phenology is simulated as a parametric function of environmental variables such as temperature and photoperiod. Parameters of these models have to be determined accurately to ensure reliable predictions.

Since model parameters often cannot be measured directly, they need to be estimated by comparing model outputs with observed data using methods such as Bayesian inference. Bayesian calibration provides a framework to quantify different sources of uncertainty, which is essential for better predictions [8], with the added value of being able to include prior information [9]. To this end, Bayesian methods have been applied in numerous crop model calibration studies [9, 10, 11, 12, 13, 14].

During the calibration of phenology models, cultivar-specific parameters are usually estimated [11]. This is because phenological traits differ markedly, not only between species and between ripening or maturity groups of crop species such as maize [15], but also between cultivars within these ripening groups. Phenological development of a cultivar is also dependent on the environment and reflects genotype × environment interactions. Thus, methods such as selecting cultivar observations from contrasting environments for calibration [13] and using cross-validation tests while evaluating environmental responses of the cultivar [16] are suggested for determining these cultivar-specific parameters.

However at regional scales, where many cultivars of a particular species are grown together, cultivar-specific parameters may not be suitable. In such calibration studies, region-specific model parameter estimates are obtained for the crop species [17, 18, 14, 19, 14], but differences between cultivars grown in the region are usually not taken into account. The resultant estimates are a compromise solution for all the cultivars grown in different environments represented by the calibration data set.

Furthermore, models may not represent the underlying processes accurately. Commonly, environmental interactions are incompletely or poorly understood, leading to conceptual uncertainty. This is reflected in multiple model formulations to represent the same process [20, 21, 22]. Consequently, models may have structural deficits. But the implicit assumption in Bayesian inference is that the model is without errors [23]. During calibration, model limitations may be compensated for by the estimated parameters [24]. As a consequence, even parameters which are meant to be cultivar-specific have been found to vary with the environment [25], thus often loosing their original physiological meaning [26].

Ignoring inherent data structures and the presence of model deficits result in inaccurate parameter estimates. When data structures such as the hierarchical classification of cultivars nested within ripening groups of a species are ignored, the uncertainty in the resultant ‘effective’ parameters are underestimated. Furthermore, indiscriminate use of large amounts of data to calibrate imperfect models leads to an overconfidence in erroneous parameter estimates [27], which in turn has been shown to result in erroneous model predictions [28]. Thus, it is important to account for these data structures and model deficits during parameter estimation.

Therefore, we propose a Bayesian multi-level calibration of a process-based plant phenology model to account for inherent data structures and to identify model deficits. Bayesian multi-level modelling (BMM) has been widely applied in ecological modelling [29, 30, 31, 32], and has more recently been applied to plant development models. For example, Patrick et al., (2009)[33] applied a hierarchical Bayesian approach to estimate parameters of the Farquhar photosynthesis model. Jarquin et al., (2016)[34] used a hierarchical Bayesian formulation of a linear-bilinear model to investigate genotype × environment (G × E) interactions of maize from breeding trials. Fer et al., (2021)[35] applied hierarchical Bayes to a dynamic vegetation model in conjunction with a Bayesian model emulator. Senf et al., (2017)[36] applied Bayesian hierarchical modelling to a satellite-based data-driven phenology model to account for spatial and temporal variation in phenology. Qiu et al., (2020)[37] developed a Bayesian hierarchical space-time model to study the impact of climate change and extreme events on phenological development. To the best of our knowledge, the BMM approach has not been applied to calibrate a process-based phenology model on a regional scale. By applying the BMM approach, we can honour the hierarchical classification of cultivars nested within ripening groups of a crop species. Thus, species-, ripening group-, and cultivar-specific parameters can be simultaneously estimated [38]. We can also account for phenological development that depends on additional environmental factors which are not already captured in the model equations [39]. Methods such as cross-calibration have been used to determine crop model parameters for representative crop cultivars grown in different agro-ecological sub-zones [40]. But parameter estimates from such an approach would have limited applications when the phenological development of a new cultivar belonging to a different ripening group is to be predicted. However, parameter estimates from the BMM approach can be used for such applications. The advantage of BMM lies in its borrowing strength [41], where parameter estimates for data-limited cultivars can benefit from data-rich ones. Additionally, the appropriate depiction of data groups results in a representative quantification of prediction uncertainty [42].

We tested the proposed approach by evaluating four BMM cases of increasing complexity in which we calibrated the SPASS [43, 44, 45] phenology model to observations of silage maize grown across Germany from 2009 to 2017. In the four BMM cases we accounted for different combinations of yearly variability, environmental effects arising from growth in different ecological regions and weather conditions, and the classification of cultivars into ripening groups. With these cases we assessed the importance of including cultivar information in regional calibration studies. We evaluated the BMM approach by comparing the calibration results from the four cases with the commonly used *pooled* approach, where a set of model parameters was estimated for silage maize grown across all environments. We also analysed trends between environmental effect parameters and environmental variables to identify possible model deficits. Thus, the findings of our study could have implications for regional calibration and model improvement studies.

## 2 Materials and Methods

### 2.1 Data

We used phenology observations of silage maize grown between 2009 and 2017 at locations across Germany, collected by the German National Meteorological Service (Deutsche Wetterdienst-DWD) [46]. The observers reported the date of the first detected occurrence of maize phenological development stages, namely, 10, 31, 53, 61, 75, 83, and 87 on the BBCH scale (Biologische Bundesanstalt, Bundessortenamt und CHemische Industrie) [47], in the assigned observation area. Corresponding cultivars and ripening groups were also reported. We refer to data sets by the site and year in which silage maize was cultivated, that is, “site-year”. We removed site-years for which sowing and harvest dates were not reported and in which the BBCH data was not strictly monotonically increasing with time. Additionally, observations that fell outside the range of sowing and harvest dates were discarded. We note that not all of the seven above-mentioned phenological stages were available in all site-years.

Minimum and maximum daily air temperatures were used as inputs to the SPASS phenology model. Weather data from the DWD stations were not available at all plant observation sites. Therefore, temperature data were extracted at all sites from the ERA5-Land re-analysis gridded data set [48]. This data set has a spatial resolution of 0.1° × 0.1°and hourly temporal resolution. The hourly data were aggregated to daily values. We consider the re-analysis data to be a better spatial representation than the point measurements at the weather stations.

To assess model limitations related to temperature and precipitation, site-years were classified into ten weather classes based on average temperature and cumulative precipitation between April and June, and between July and September. A K-means clustering algorithm was applied to define the weather classes (details in Appendix B). Site-years were also grouped into nine ecological regions based on the classification provided by the Bundesamt für Naturschutz [49]. For computational reasons, a subset of 100 site-years out of 3,004 was randomly selected for calibration where it was ensured that at least one site-year was selected from each of the four ripening groups, nine ecological regions, ten weather classes and nine years. The calibration data set consisted of 66 cultivars from the four ripening groups (Table 1). It was also ensured that the relative proportions of site-years from the different ripening groups in the full data set were maintained in the calibration subset (early: 34%, mid-early: 54%, mid-late: 11% and late: 0.4% in the full data set).

**Table 1:** Summary of site-years and cultivars used for calibration

| ripening group | early | mid-early | mid-late | late | Total |
| --- | --- | --- | --- | --- | --- |
| number of cultivars | 25 | 33 | 7 | 1 | 66 |
| number of site-years | 35 | 55 | 9 | 1 | 100 |

### 2.2 Phenology model

Air temperature, site-latitude, and sowing and harvest dates are required as inputs to the SPASS phenology model. The model has nine parameters, seven of which were estimated during calibration (Table 2) and the remaining two were fixed at default values. The model equations and details are given in Appendix A. We provide a brief summary below.

**Table 2:**
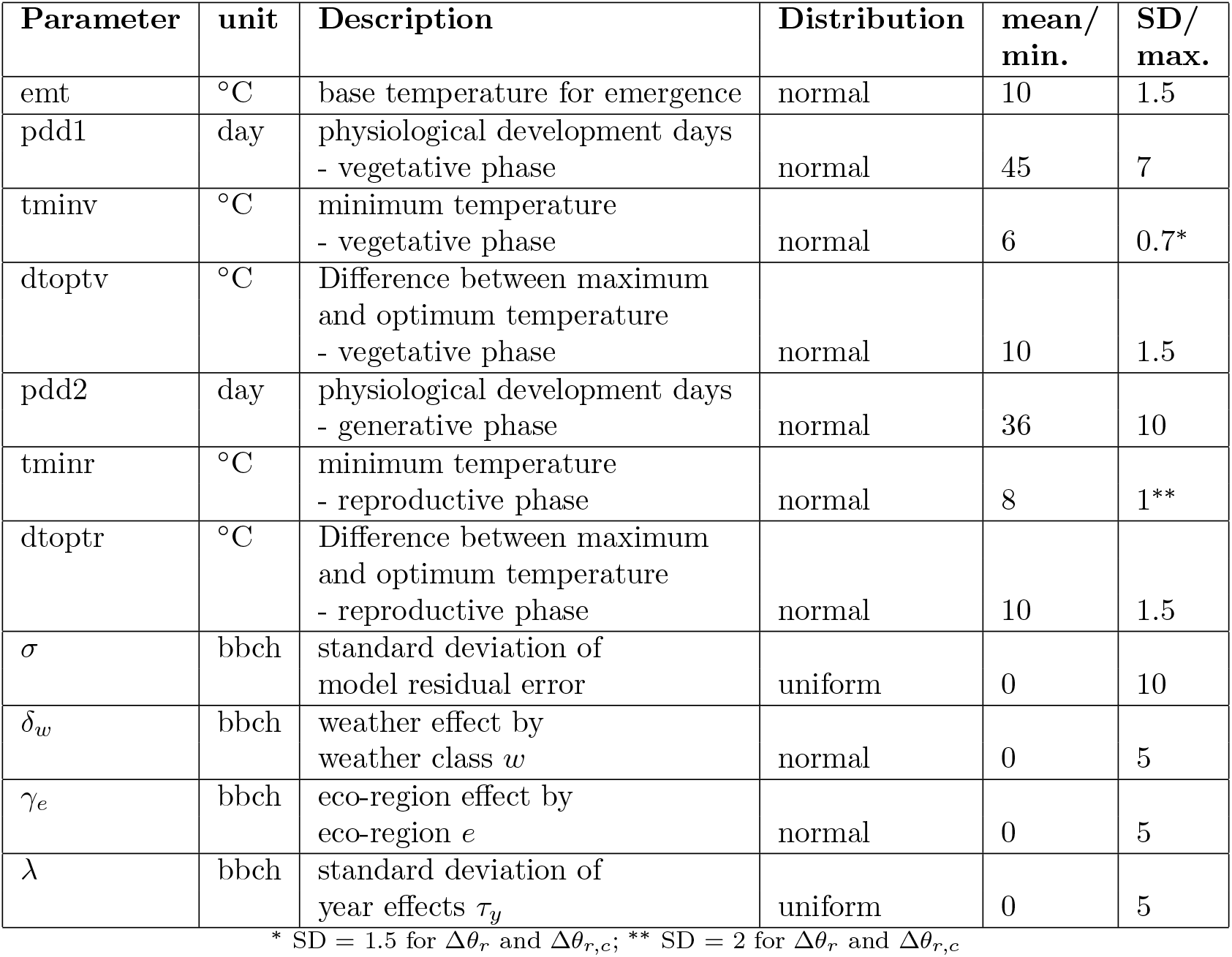
Prior distributions for estimated parameters in the process-based SPASS phenology model and the multi-level model cases. The mean and standard deviation (SD) are specified for normal distributions while the minimum (min.) and maximum (max.) are specified for uniform distributions.

Three main phases of development are defined in the model: emergence, vegetative and reproductive growth phases. Emergence is dependent on the sowing depth, assumed to be fixed for all the site-years at 3 cm, and on the temperature above a minimum value (*emt*). The development rate during the vegetative and reproductive phases is dependent on the number of physiological development days at optimum temperature (*pdd1* for vegetative and *pdd2* for reproductive) and on the Temperature Response Function (TRF). The TRF is defined by phase-specific minimum, optimum and maximum cardinal temperatures (*tminv, toptv* and *tmaxv*, respectively, for vegetative and *tminr, toptr* and *tmaxr*, respectively for reproductive). The SPASS phenology model as described in Wang (1997)[43] was implemented in our study with the following modifications: (a) the photoperiod effect on the vegetative phase was not considered, (b) no soil water-limiting effect on germination was assumed and germination occurs instantaneously after sowing, and (c) for numerical reasons the transition between emergence and vegetative phases was defined by a sigmoidal function instead of the original step function.

The parameters for physiological development days at optimum temperature for the vegetative (*pdd1*) and reproductive phases (*pdd2*) were estimated in the study. Additionally, minimum and optimum temperature for vegetative (*tminv* and *toptv*, where *toptv* = *tmaxv–dtoptv*) and reproductive (*tminr* and *toptr*, where *toptr* = *tmaxr* – *dtoptr*) phases, as well as the minimum temperature required for emergence (*emt*), were estimated. However, the parameters *tmaxv* and *tmaxr* were not estimated. The range of average daily temperatures during the growing season at the study site-years were between −6 and 31°C. This is usually expected to be at or lower than the optimal temperatures for maize, a warm-weather plant. The lack of observations in the supra-optimal temperature range would make constraining *tmax* difficult [50] and is expected to incur problems of equifinality. To avoid these problems, the values of *tmaxv* and *tmaxr* were fixed at 44° C.

We used Bayesian inference to determine the posterior probability of the model parameters. Let *ϕ_d_* represent the given phenology observation on day *d*. The phenology 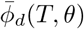 at day *d*, simulated by the SPASS model with a parameter vector *θ*, is dependent on air temperatures *T* from the date of germination to the day *d*. The phenology observations are available at days *D*, so the parameters are conditioned on Φ = {*ϕ_d_*; *d* ∈ *D*} through the likelihood function *p*(Φ | *θ,T*). The posterior parameter distribution is given by

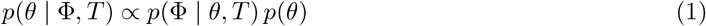

where *p*(*θ*) is the joint prior probability distribution of the parameters.

### 2.3 Bayesian model cases

We describe five Bayesian model cases (one pooled and four multi-level model cases as seen in Fig.1) in terms of their likelihood functions in the following sections. The prior distributions for all the estimated parameters are provided in Table 2.

**Figure 1:**
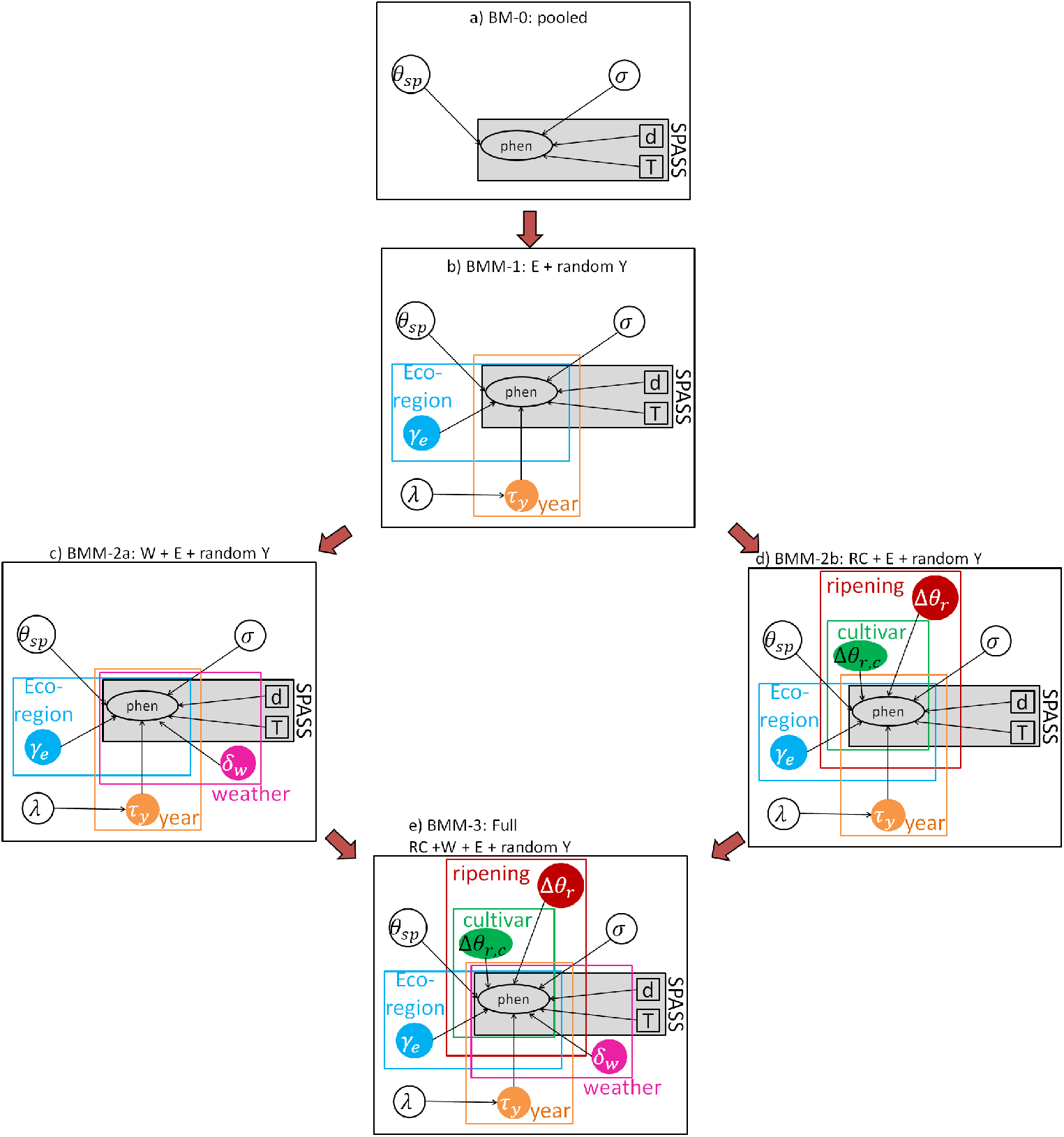
Graphical representation of the five Bayesian models: (a) BM-0: pooled, (b) BMM-1: ecoregion effects with random year effects, (c) BMM-2a: weather and eco-regions effects with random year effects, (d) BMM-2b: hierarchical classification of cultivars into ripening groups with eco-regions effects and random year effects, and (e) BMM-3 or full model: hierarchical classification of cultivars into ripening groups, eco-region and weather effects and random year effects. In the SPASS phenology model, phenological development on a given day (*d*) is a function of air temperatures (*T*) from the date of germination to that day. *θ_sp_* is the maize species-level parameter vector, Δ*θ_r_* is the difference between the ripening group-level parameter *θ_sp,r_* and *θ_sp_*, Δ*θ_r,c_* is the difference between the cultivar-level parameter *θ_sp,r,c_* and *θ_sp,r_*, and *σ* is the standard deviation of the likelihood function. *γ_e_* represents the eco-region effect, *δ_w_*, the weather class effect, *τ_y_* the year effect and λ is the standard deviation of the year effect. E=eco-regions, Y=year,W=weather class, R=ripening group, C=cultivar. The red arrows outside the model sketches represent model extensions. During calibration, *θ_sp,r_* is estimated for the 4 ripening groups, *θ_sp,r,c_* for 66 cultivars, *δ_w_* for 10 weather classes, *γ_e_* for 9 eco-regions, and *τ_y_* for 9 years.

#### 2.3.1 BM-0: Pooled model

The pooled model is the most commonly used calibration setup for regional scale studies, where a common parameter set is estimated for all cultivars grown in different environmental conditions in the region. Assuming independent Gaussian observation errors, the likelihood function for the pooled model is given by

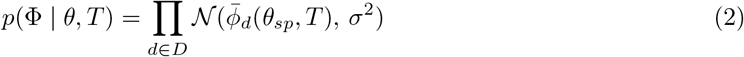

where *θ* = {*θ_sp_, σ*}, *θ_sp_* represents the maize species (*sp*) parameters and 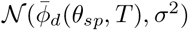 the density of a normal distribution with mean equal to the simulated phenology 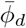 and standard deviation *σ*. The pooled model case is shown in Fig. 1a. The joint prior probability 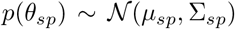 was represented by a multivariate normal distribution with seven dimensions corresponding to the SPASS model parameters (*emt, pdd1, tminv, dtoptv, pdd2, tminr, dtoptr*). The mean vector of the distribution (*μ_sp_*) and main diagonal elements of the variance-covariance matrix (Σ_*sp*_) are defined in Table 2 (mean and squared standard deviation, respectively) while the off-diagonal elements are zero.

#### 2.3.2 BMM-1: Fixed eco-region effects and random year effects

We expect that the different eco-regions and years in which silage maize was grown in Germany influence phenology. We analysed this effect with the BMM-1 model (Fig. 1b), where we accounted for fixed effects due to the different eco-regions and random effects arising from variability between the years. If the different eco-regions and years are represented by *e* ∈ *E* and *y* ∈ *Y*, respectively, then

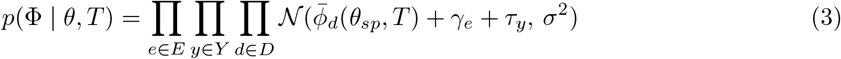

where parameters *γ_e_* and 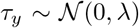 represent the effects by eco-region *e* and year *y*, respectively, *E* = 9 is the total number of eco-regions, *Y* = 9 is the total number of years, and *θ* = {*θ_sp_, γ_e_*, λ, *σ*}. A uniform prior density was assumed for the standard deviation of the year effects λ as per Gelman (2006) [51] (Table 2).

#### 2.3.3 BMM-2a: Fixed eco-region and weather effects and random year effects

Although the SPASS model accounts for the effect of temperature on phenological development, there could be other weather conditions (i.e. temperature and precipitation during specific phases) that are important but not adequately captured in the model. Also, differences in weather conditions could result in a perceived variability between eco-regions and between years. In the BMM-2a model (Fig. 1c), we additionally accounted for the effects due to the different weather classes. If the different weather classes are represented by *w* ∈ *W* and parameter *δ_w_* represents the effects by weather class, then

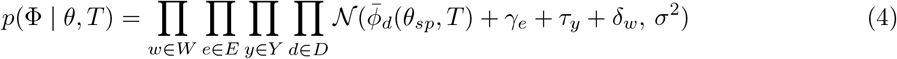

where 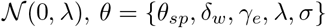 and *W* = 10 is the total number of weather classes.

#### 2.3.4 BMM-2b: Fixed ripening, cultivar, eco-region effects and random year effects

As a modification from BMM-1, we also accounted for the inherent structure in the data in BMM-2b (Fig. 1d) wherein the cultivars c, are nested within ripening groups *r* of the maize species *sp*.

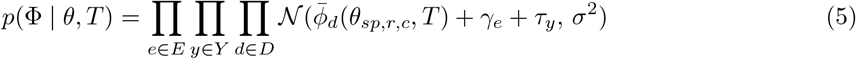

where *θ* = {*θ_sprc_, γ_e_*, λ, *σ*} and *θ_sp,r,c_* represents the joint probability distribution of all the estimated cultivar-level parameters in the hierarchy. It can be expressed as *θ_sp,r,c_* = *θ_sp_* + Δ*θ_r_* + Δ*θ_r,c_*, where Δ*θ_r_* = *θ_sp,r_* – *θ_sp_* is the difference between the species-level parameters (*θ_sp_*) and the ripening group-level parameters (*θ_sp,r_*), and Δ*θ_r,c_* = *θ_sp,r,c_* – *θ_sp,r_* is the difference between the ripening group-level and the cultivar-level parameters. Thus, the SPASS model parameters corresponding to 66 cultivars (cultivar-level) and 4 ripening groups (ripening group-level) in the calibration data set are estimated. Their prior probability 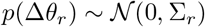 and 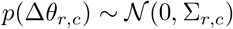 were represented by multivariate normal distributions, centred at zero. Their variance-covariance matrices (Σ_*r*_, Σ_*r,c*_) were equivalent to that of the species-level prior Σ_*sp*_ for all parameters except *tminv* and *tminr* (note different standard deviation in the footnote of Table 2).

#### 2.3.5 BMM-3: Full model

Finally, in the full model (Fig. 1e), we accounted for the inherent hierarchical data structure, ecoregions and weather effects, as well as year effects.

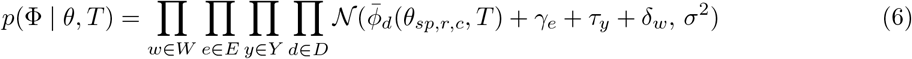

where *θ* – {*θ_sp,r,c_, δ_w_, γ_e_*· *σ*} and *θ_sp,r,c_* – *θ_sp_* + Δ*θ_r_* + Δ*θ_r,c_*.

### 2.4 Posterior sampling

Markov Chain Monte Carlo sampling of the posterior parameter distributions was performed using the Gibbs algorithm from the Jags software [52] implementation in R2jags [53] and jagsUI [54] packages in R [55]. For the model cases BM-0, BMM-1, BMM-2a and BMM-2b, 500 runs were used for adaptation. Three chains were run and 5000 iterations were run per chain until the Gelman Rubin convergence diagnostic was ≤ 1.1. Of these iterations, every 5th parameter vector (thinning = 5) was stored, resulting in a total of 3,000 samples that were used for generating the posterior parameter distributions and simulated phenology described in the results. For BMM-3, 100 runs were used for adaptation. Three chains were run and 3,600 iterations per chain were run until the Gelman Rubin convergence diagnostic was ≤ 1.1. All the samples (total of 10,800) were used for the plots. Diagnostic plots for the MCMC samples are provided in Supplement S1.

### 2.5 Calibration performance metrics

Calibration quality was assessed for each of the 100 site-years used for calibration and for each of the five model cases by estimating the expected value 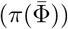 of a loss function 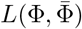 where 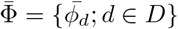 is a vector of phenological development simulated by the model.

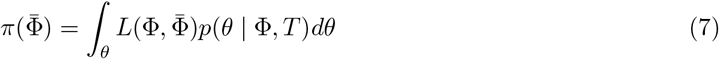

where 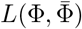 is either 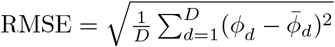 or 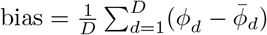.

## 3 Results

We first provide results from the classification of site-years into environmental classes (section 3.1). We then describe the calibration quality of the SPASS model in the different Bayesian model cases (section 3.2), followed by an analysis of the posterior distributions of the SPASS model parameters (section 3.3), the environmental effects (section 3.4) and residual uncertainty (section 3.5). All figures were made using the ggplot2 [56] package in R.

### 3.1 Classification of site-years into environmental classes

All site-years were classified into ten weather classes (Fig. 2). The weather classes were based on the average temperatures and cumulative precipitation between April and June, and between July and September. Silage maize cultivated across Germany generally undergoes vegetative development from April to June and reproductive development from July to September. Phenological development during the vegetative and reproductive phases are dependent on temperature. The relationship between temperature and phenological development is usually represented by equations in phenological models, including SPASS. However, existing model equations may not accurately capture this temperature response. Additionally, the influence of factors like precipitation that are known to influence phenology in some plant species [57], could also have either a direct influence on maize phenology or an indirect effect by influencing temperatures within the crop canopy. However, these effects are not represented in the SPASS phenology model. Thus, site-years were classified into the weather classes to assess model limitations related to temperature and precipitation. All site-years were also classified into nine ecological regions (Fig. 3). Note that since the 100 site-years used for calibration were randomly sampled, the calibration data set contained only a few site-years from ecological regions in the northeastern part of Germany.

**Figure 2:**
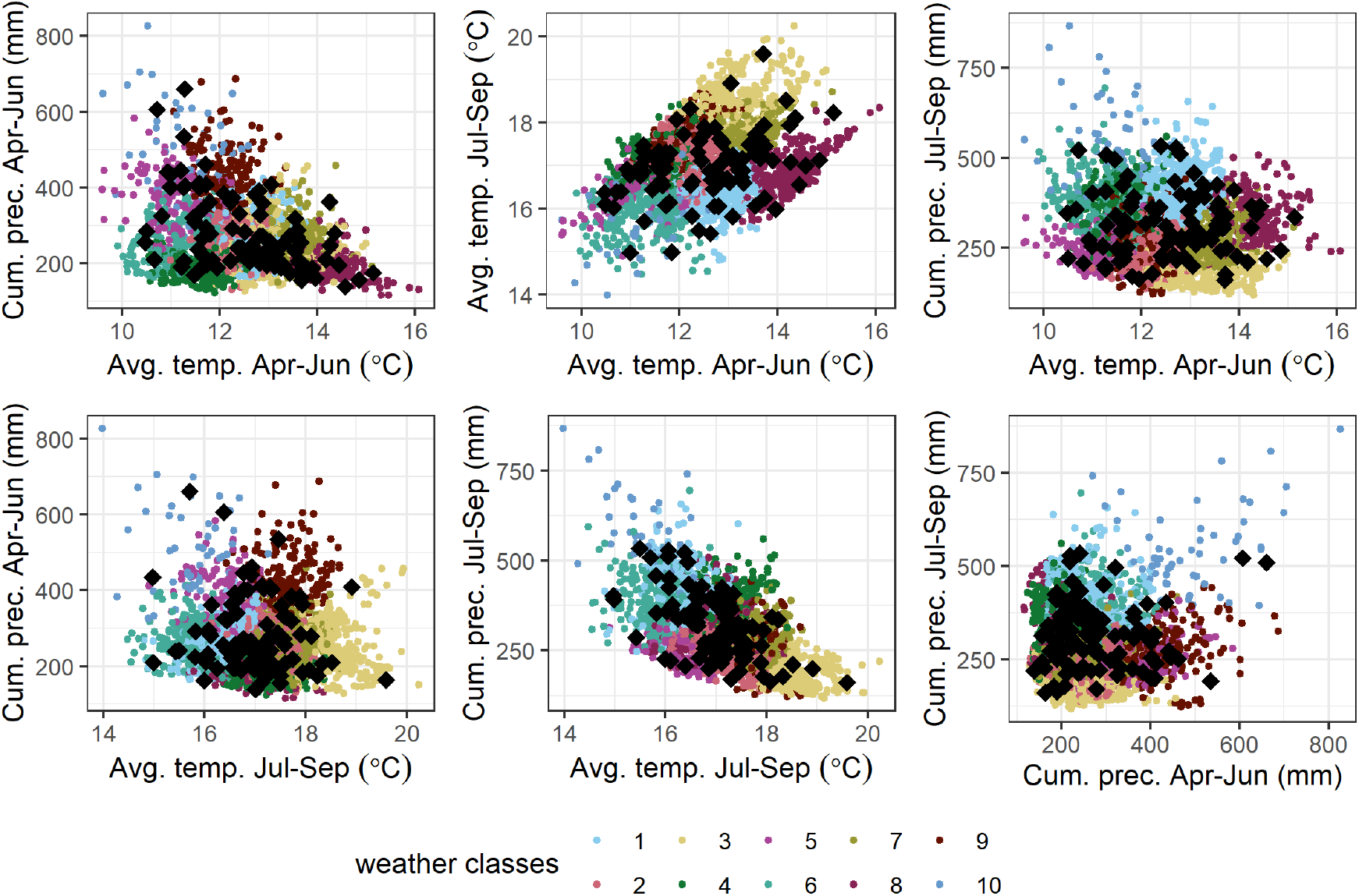
Site-years of silage maize grown across Germany were classified into ten weather classes based on average temperature (avg. temp.) and cumulative precipitation (cum. prec.) during April-June (Apr-Jun) and July-September (Jul-Sep). Black diamonds are the 100 site-year samples selected for calibration from the overall data set of site-years indicated by the coloured points. The colours indicate the ten identified weather classes.

**Figure 3:**
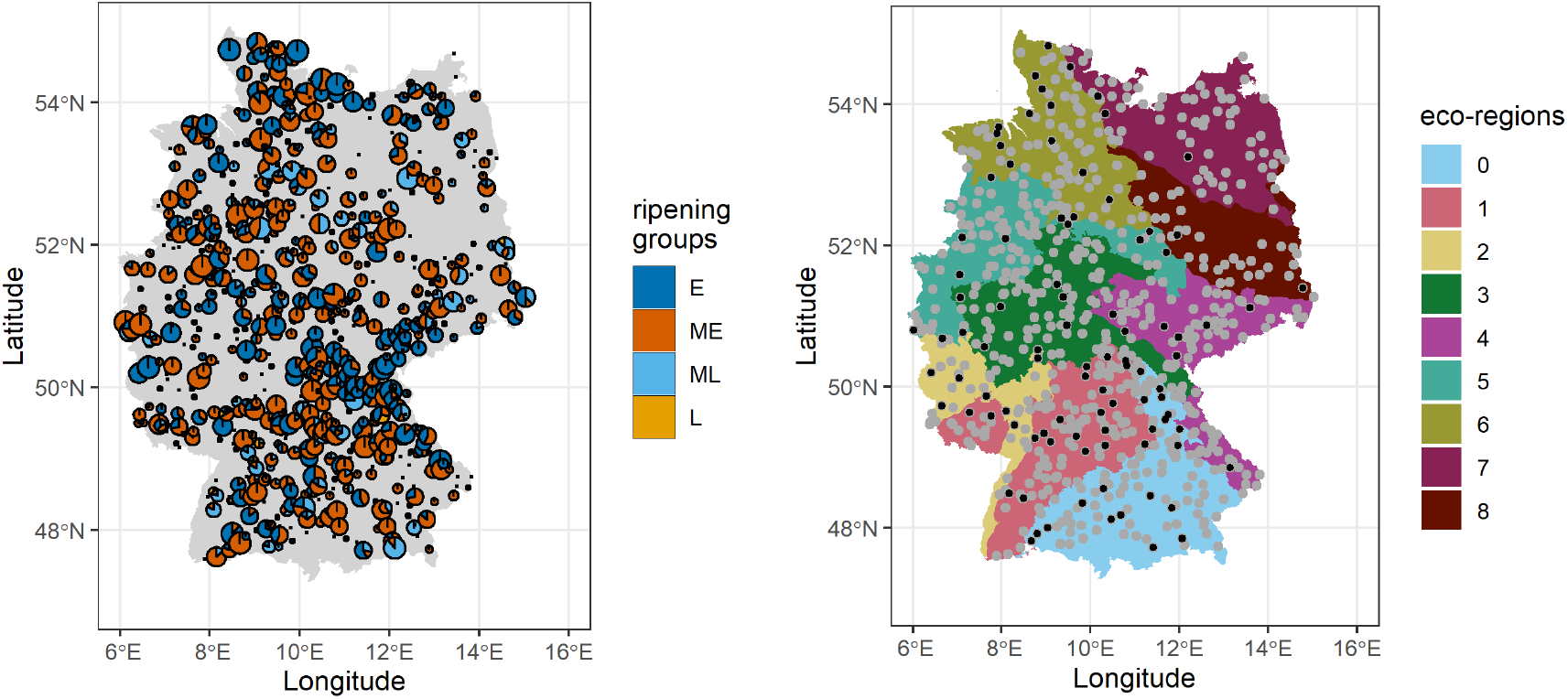
Ripening groups of silage maize cultivated in nine eco-regions across Germany from 2009-2017. (a) Pie-charts of silage maize ripening groups grown at different sites in Germany. (b) Location of different sites (points) where silage maize was grown across the nine eco-regions (coloured polygons). The black points are the site locations of the 100 site-year samples used for calibration. (Projection system: DHDN 3 Degree Gauss Zone 3)

### 3.2 Calibration quality

As an example, we compare observed and simulated phenology for silage maize grown at a site in the state of Bavaria in 2009 from the pooled model case (BM-0) and the full model case (BMM-3) (Fig. 4). The blue bands show the 5-95^th^ percentile of simulated phenology that accounts for uncertainty from model parameters (*θ_sp_* for BM-0 in Fig. 4a and *θ_sp,r,c_* for BMM-3 in Fig. 4b) while the red bands additionally accounts for environmental effects (*δ_w_, γ_e_*, *τ_y_* in Fig. 4b). The grey bands show the 5-95^th^ percentile of simulated phenology that additionally accounts for the unresolved residual error (*σ*). There is a reduction in bias in BMM-3 as compared to BM-0. Furthermore, there is an overall reduction in unresolved residual error in BMM-3 with the model parameters and environmental effects accounting for a large share of the error variance. We also note that the blue bands in Fig. 4(a) have collapsed around the mean simulated phenology.

**Figure 4:**
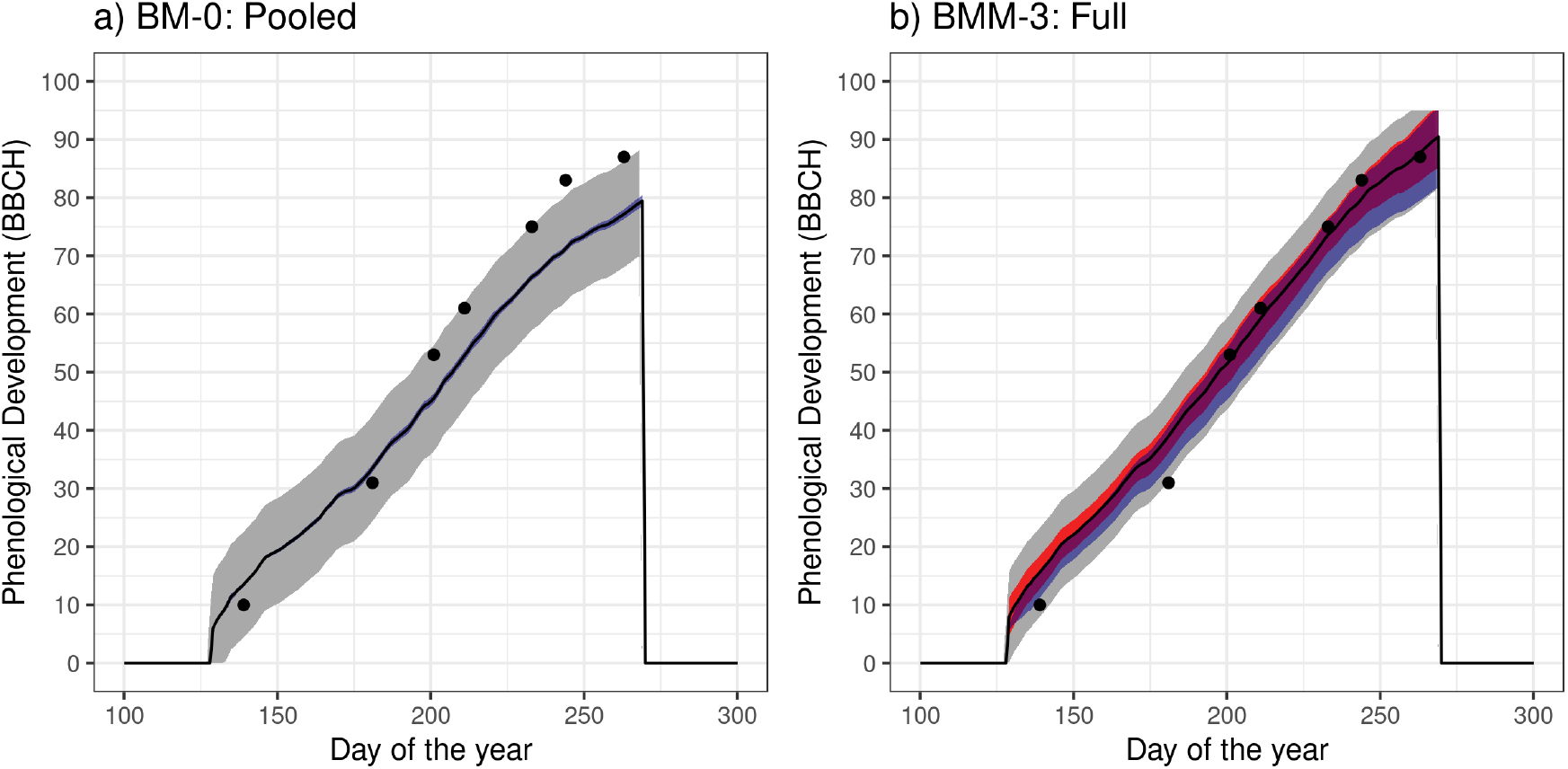
Observed and simulated phenology of silage maize (cultivar: Amatus) at Benediktbeuern in the state of Bavaria in 2009 in (a) the pooled model case (BM-0) and (b) full model case (BMM-3). The black points are the observations. The solid black line is the mean simulated phenology, while the coloured bands represent 5-95^th^ percentile of simulated phenology: The blue bands represent the uncertainty originating from process model parameters, while the red band in (b) represents the uncertainty originating from process model parameters and environmental effects (weather, eco-region and year). The grey bands additionally account for the unresolved residual error.

The model performance represented by the mean RMSE and bias for the 100 calibration site-years (Fig. 5) improved with model complexity from the pooled model to the full model (BM-0, BMM-1, BMM-2a, BMM-2b, BMM-3). This is evident from the reduction in mean RMSE and shrinkage of the mean bias towards zero. In Fig. 6 the model performance from the five cases were analysed by ripening group and weather class. Across the plots, the two cases BMM-2b and BMM-3 that account for the ripening group-cultivar hierarchy generally exhibit a lower bias and RMSE as compared to the others. Across the four ripening groups (Fig. 6a), the mean bias is closer to zero with increasing model complexity, as seen in Fig. 5. A decrease in mean bias and RMSE occurs on the inclusion of cultivar-ripening group information through the hierarchy in BMM-2b and BMM-3. The single site-year from the late ripening cultivar included in calibration also exhibits a clear improvement in RMSE and bias. While the inclusion of weather effects (Fig. 6b) in the model cases BMM-2a and BMM-3 result in smaller mean RMSE and bias only in some weather classes, the inclusion of cultivar-ripening group hierarchy results in an improvement in most classes. Although the inclusion of eco-regions and year effects (BMM-1 to BMM-3) (Fig. C.12(a) and (b) in the Appendix C) improves RMSE and bias in some eco-regions and years, a clear trend across all the classes cannot be identified.

**Figure 5:**
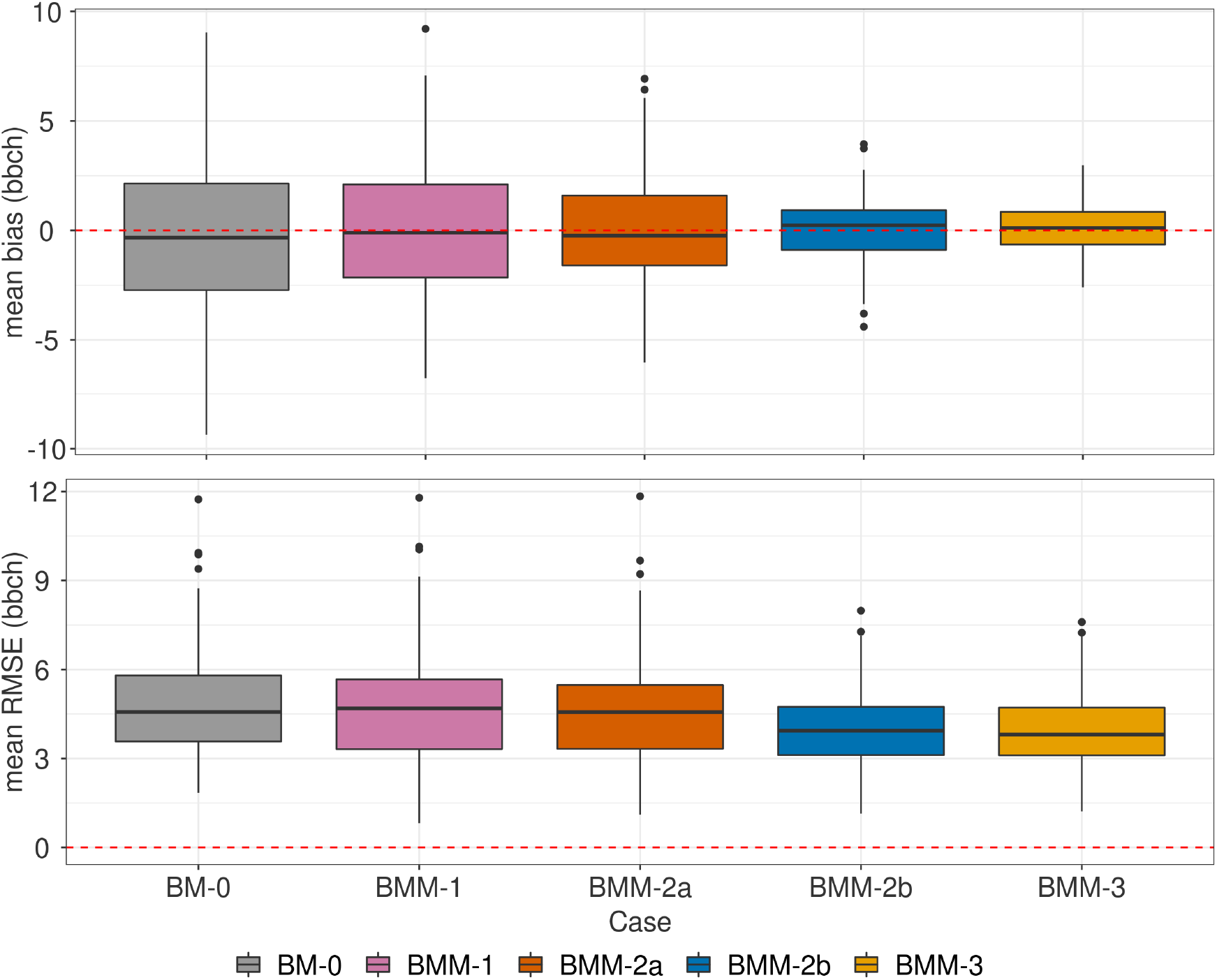
Box-plots of the mean RMSE and bias for each calibration site-year in the five model cases, BM-0 (pooled), BMM-1 (eco-regions, random year effects), BMM-2a (eco-regions, weather class, random year effects), BMM-2b (ripening group-cultivar hierarchy, eco-region, random year effects), BMM-3 (full model). Each box-plot represents the 100 site-years used for calibration. Hinges of the box-plot represent the inter-quartile range (IQR), whiskers extend from the hinges up to 1.5×IQR and values beyond this range are plotted as points.

**Figure 6:**
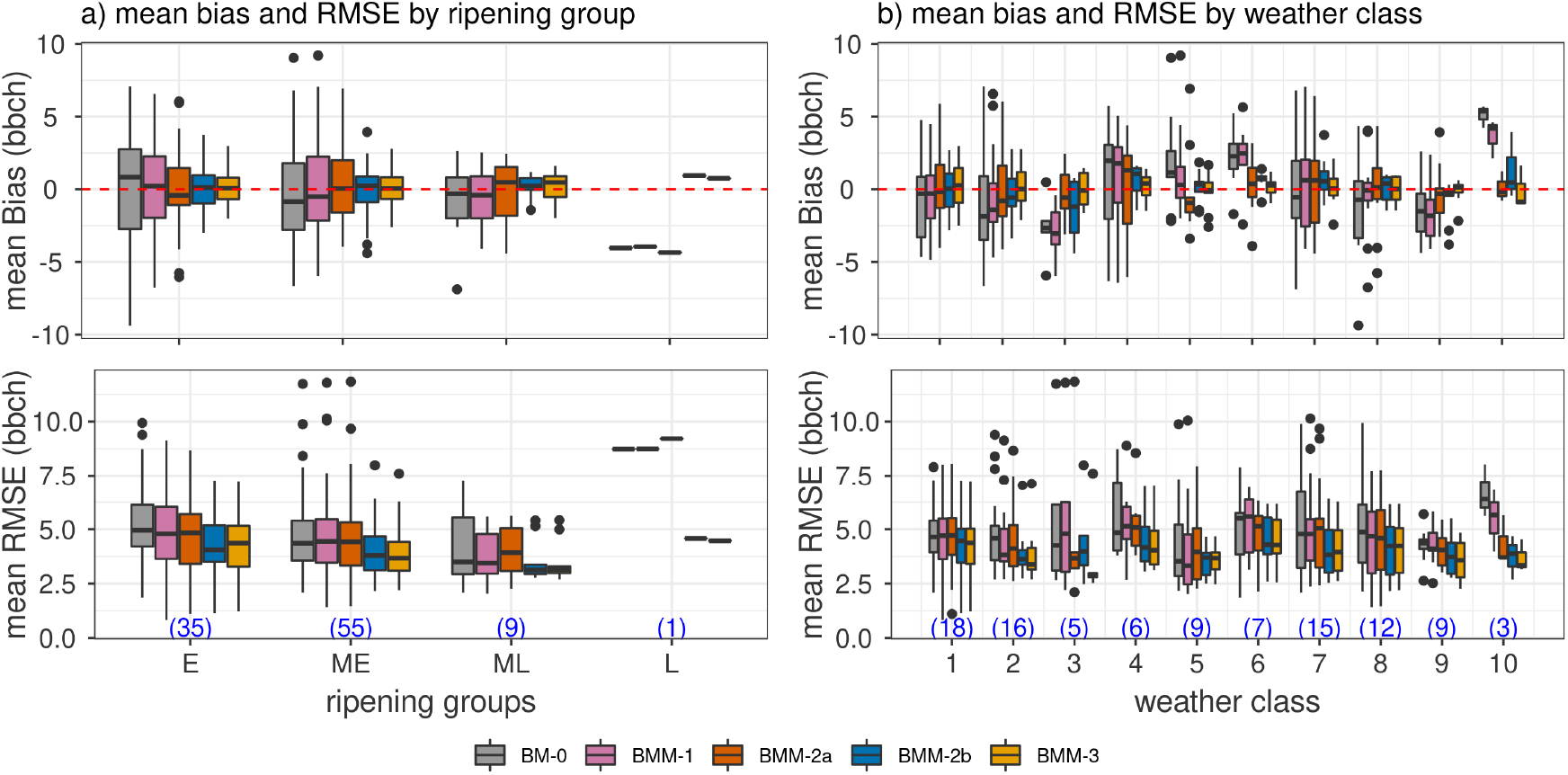
Box-plots of the mean RMSE and bias by site-year within ripening groups (a), weather class (b) for each of the five model cases, BM-0 (pooled), BMM-1 (eco-regions, random year effects), BMM-2a (eco-regions, weather class, random year effects), BMM-2b (ripening group-cultivar hierarchy, eco-region, random year effects), BMM-3 (full model). The numbers in blue at the bottom of the two subplots indicate the number of site-years and consequently the number of points in the groups defined on the x-axis that were used to obtain the box-plots. Hinges of the box-plot represent the inter-quartile range (IQR), whiskers extend from the hinges up to 1.5×IQR and values beyond this range are plotted as points.

### 3.3 Phenology model parameters

The marginal posterior distributions of the SPASS model parameters were analysed for the full model (BMM-3) to investigate differences between cultivar, ripening group and maize species parameter estimates after the environmental effects are taken into account. Figure 7 shows the posterior parameter distribution by species (*θ_sp_*), ripening groups (*θ_sp,r_* = *θ_sp_* + Δ*θ_r_*) and cultivars (*θ_sp,r,c_* = *θ_sp_* + Δ*θ_r_* + Δ*θ_r,c_*) for two parameters *tminv* and *pdd1.* Parameter *tminv* shows low variability between cultivars of the same ripening group while *pdd1* shows high variability. A similar visual inspection of other parameters showed that they could be classified into the categories of low (*tminv, tminr, toptr*) and high (*pdd1, pdd2, toptv, emt*) between-cultivar variability (Fig.D.13 and D.14 in Appendix D).

**Figure 7:**
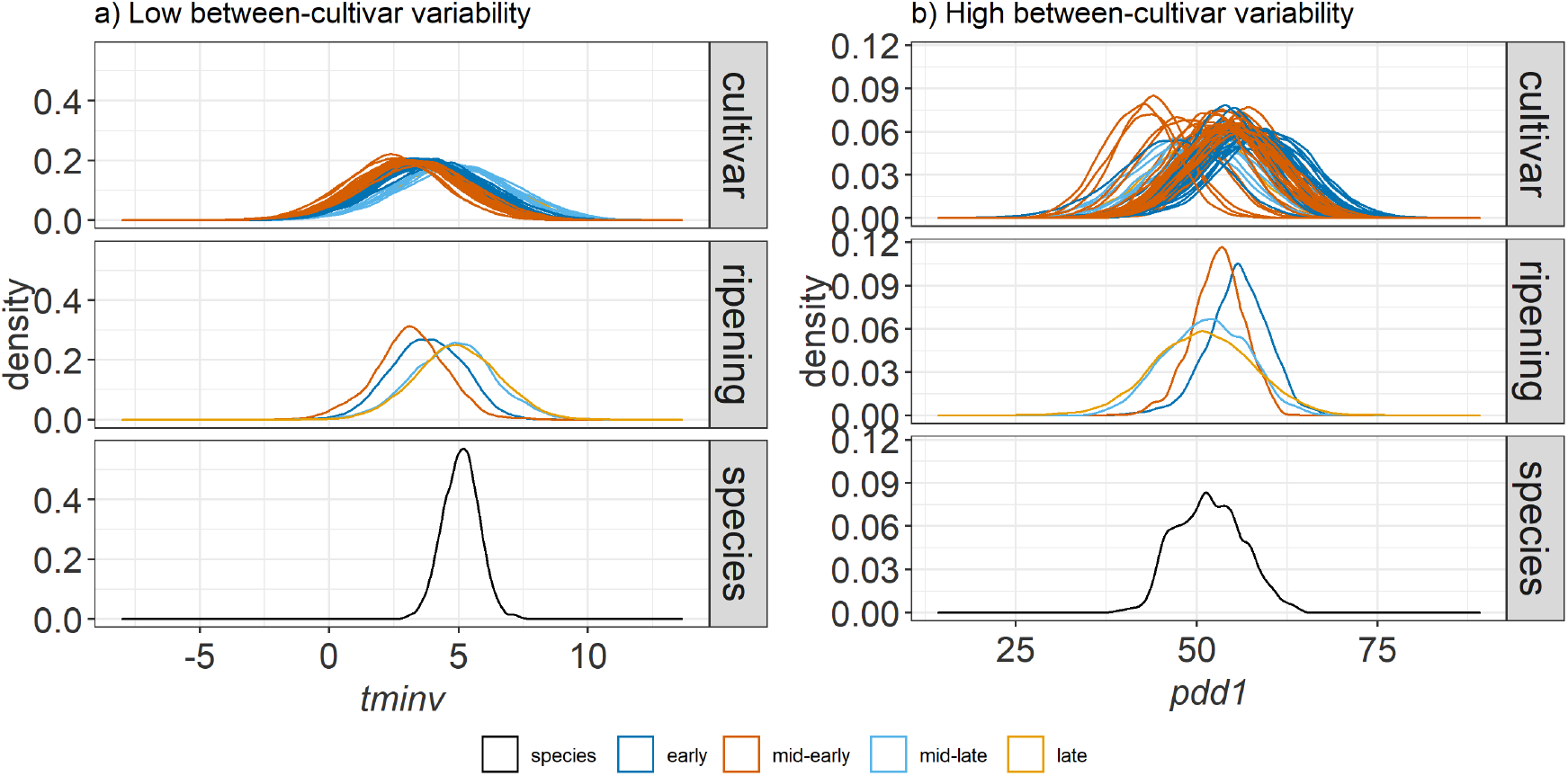
Posterior distribution of parameters in the full model case BMM-3: minimum temperature for development in the vegetative phase (*tminv*) and physiological development days for the vegetative phase at optimum temperature (*pdd1*). The distributions are provided for the species (*θ_sp_*), ripening group (*θ_sp,r_*) and cultivar (*θ_sp,r,c_*) levels of the hierarchy. Colours of cultivar distributions correspond to their respective ripening groups.

### 3.4 Environmental effects

The prior and posterior parameter distributions of the weather effects (*δ_w_*) and eco-region effects (*γ_e_*) were analysed with respect to their corresponding classes in the four multi-level model cases (Fig. 8). The parameter distributions of the weather effects are only shown for BMM-2a and BMM-3 since these effects are taken into account only in these two cases. The posterior parameter distributions deviate from the prior which is normally distributed around zero and are narrower than the prior.

**Figure 8:**
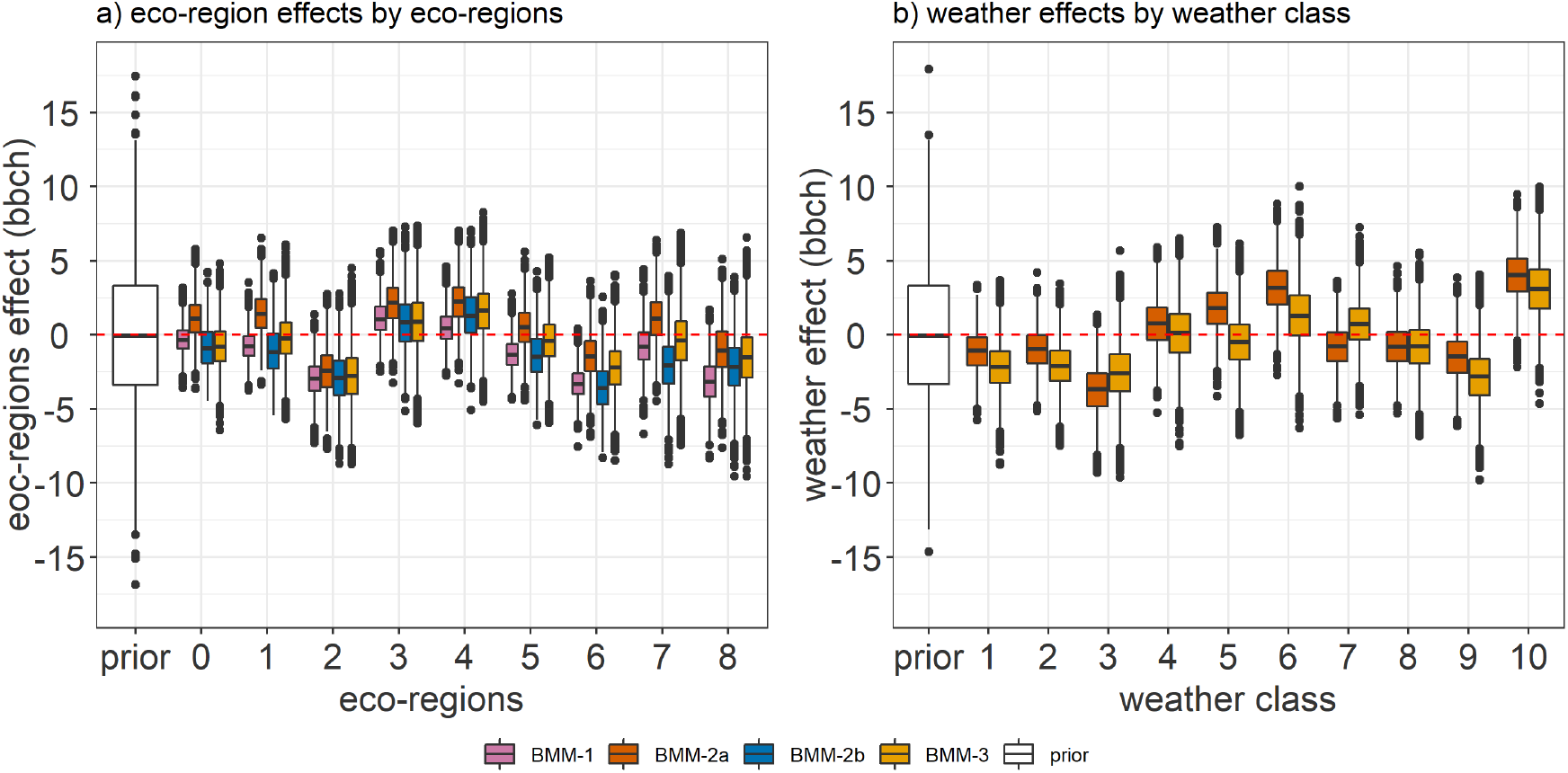
The prior and posterior parameter distributions of (a) eco-region effects for the nine ecoregions and (b) weather effects for the ten weather classes. The distributions of the eco-region effects (*γ_e_*) and weather effects (*δ_w_*) (y-axis) are plotted against their corresponding classes (x-axis) in the four Bayesian multi-level model cases. The Bayesian multi-level models are: BMM-1: ecoregion with random year effects; BMM-2a: weather, eco-region with random year effects; BMM-2b: cultivar-ripening group hierarchy, eco-region and random year effects; BMM-3: cultivar-ripening group hierarchy, weather, eco-region random year effects. Hinges of the box-plot represent the inter-quartile range (IQR), whiskers extend from the hinges up to 1.5×IQR and values beyond this range are plotted as points.

To identify possible model deficits, we analysed trends between the median value of the weather effects parameters and mean of the average daily temperature and cumulative precipitation of the weather classes from April to June and July to September for the cases BMM-2a and BMM-3. A high correlation coefficient is seen between the median weather effect per class and the mean of the average daily temperature from July to September (Fig.9). The correlation coefficient reduces from −0.87 in BMM-2a to −0.64 in the full model BMM-3 where the cultivar-ripening group hierarchy is considered. This is also accompanied by a widening in confidence intervals of the linear regression line.

**Figure 9:**
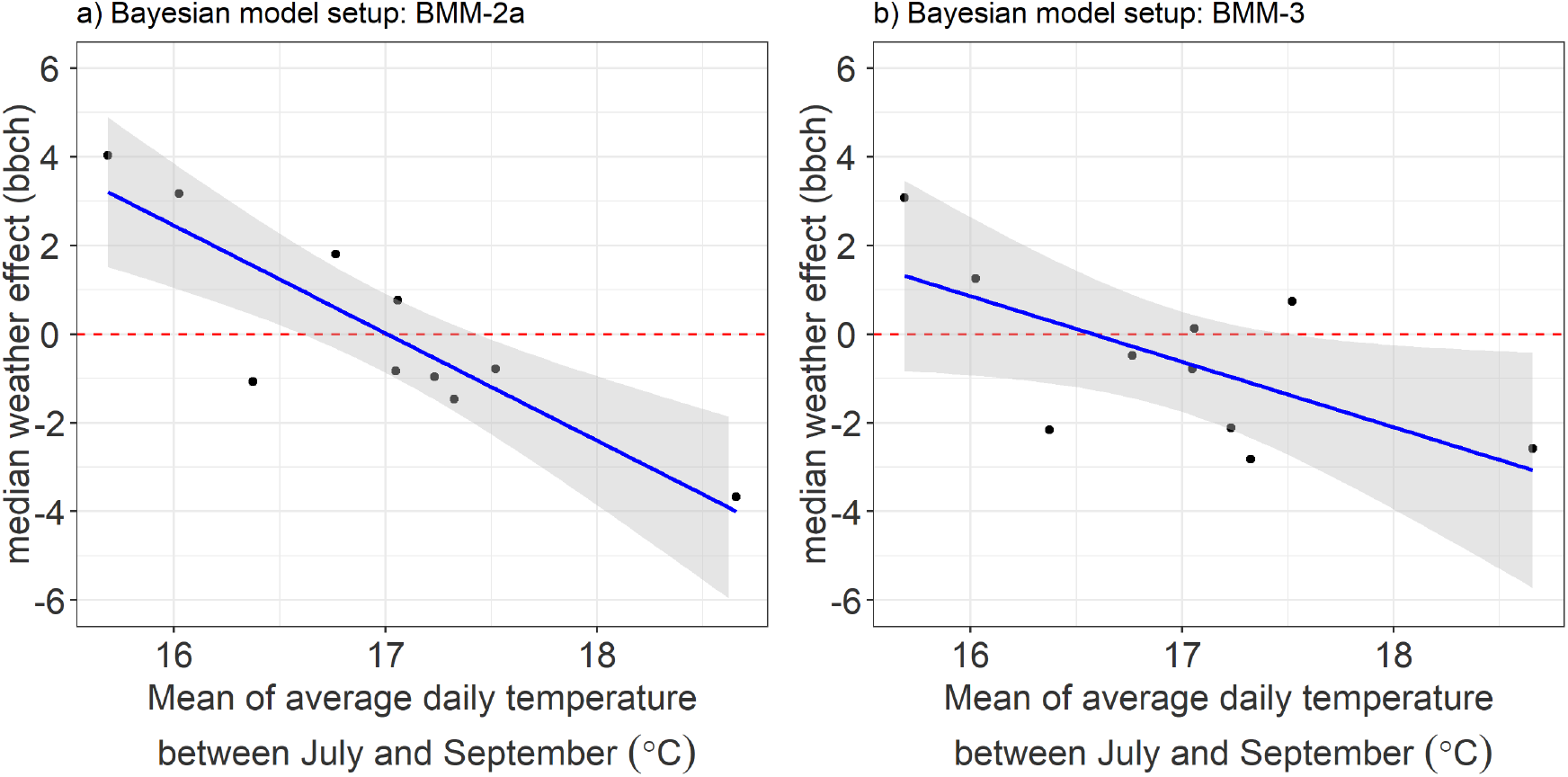
Median value of the weather effects parameters plotted against the mean of the average daily temperature between June and September for the 10 weather classes for the Bayesian model case (a)BMM-2a and (b)BMM-3. The ten points correspond to the ten weather classes. The grey bands represent 95% confidence interval of the regression line.

The eco-region effect (*γ_e_*) was included in all the Bayesian multi-level model cases. Figure 10 shows a comparison between the median eco-regions effects for the four multi-level models. A negative eco-region effect indicates an overestimation of phenology by the model and the other effects that were accounted for in the particular BMM case. This overestimation was corrected by the eco-region effect parameter. Conversely, a positive eco-region effect indicates an underestimation of phenology by the model and the other effects. Eco-regions 6 and 8 have similar median eco-region effects, and so do 3 and 4. Eco-regions 2, 6, and 8 show a negative eco-region effect while 3 and 4 show a positive effect irrespective of the model case. A comparison of BMM-1 (Fig. 10a) with BMM-2a (Fig. 10b) and BMM-2b (Fig. 10c) shows that the inclusion of weather effects (BMM-2a) results in a positive eco-region effect in most regions, while the inclusion of cultivar-ripening group hierarchy (BMM-2b) results in negative eco-region effects. However, this tendency is not seen when both weather effects and cultivar-ripening group hierarchy are included in BMM-3 (Fig. 10d).

**Figure 10:**
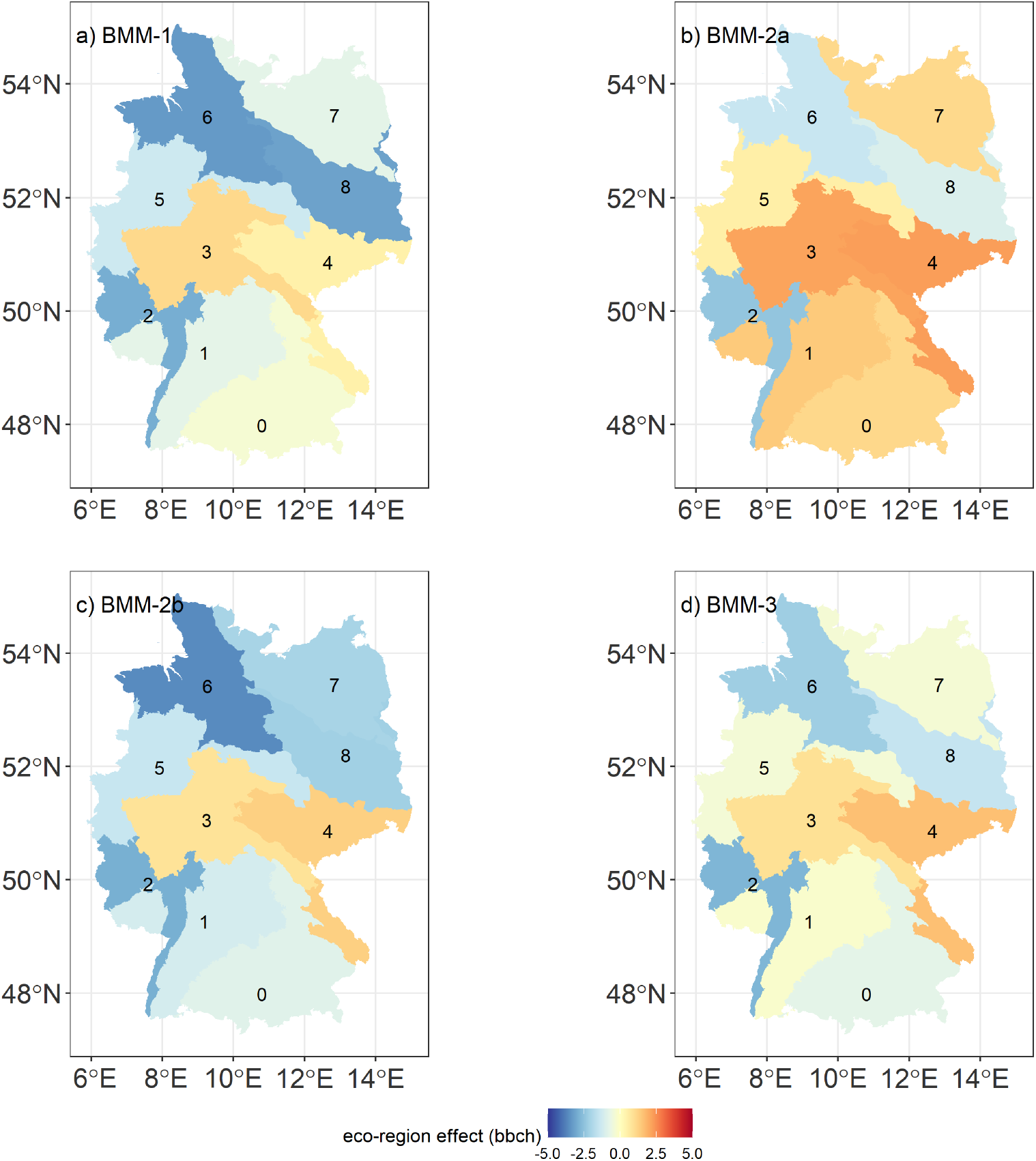
Median value of the eco-region effects parameters plotted for the nine eco-regions across Germany for the Bayesian multi-level models (a) BMM-1 (b) BMM-2a, (c) BMM-2b, and (d) BMM-3. The numbers indicate the nine eco-regions. (Projection system: DHDN 3 Degree Gauss Zone 3)

### 3.5 Residual uncertainty

The standard deviation (*σ*) of the likelihood function and standard deviation (λ) of the year effect *τ_y_*, represent the unresolved and resolved components of residual uncertainty, respectively. Figure 11a shows the posterior parameter distribution of *σ* from the five Bayesian models. There is a reduction in unresolved residual uncertainty with increasing model complexity from BMM-0 to BMM-3. A large reduction is seen on the inclusion of ripening-cultivar hierarchy in BMM-2b and BMM-3. The standard deviation of the year effect (λ) (Fig.11b) shows an increase from BMM-1, BMM-2a to BMM-2b, followed by a slight decrease in BMM-3.

**Figure 11:**
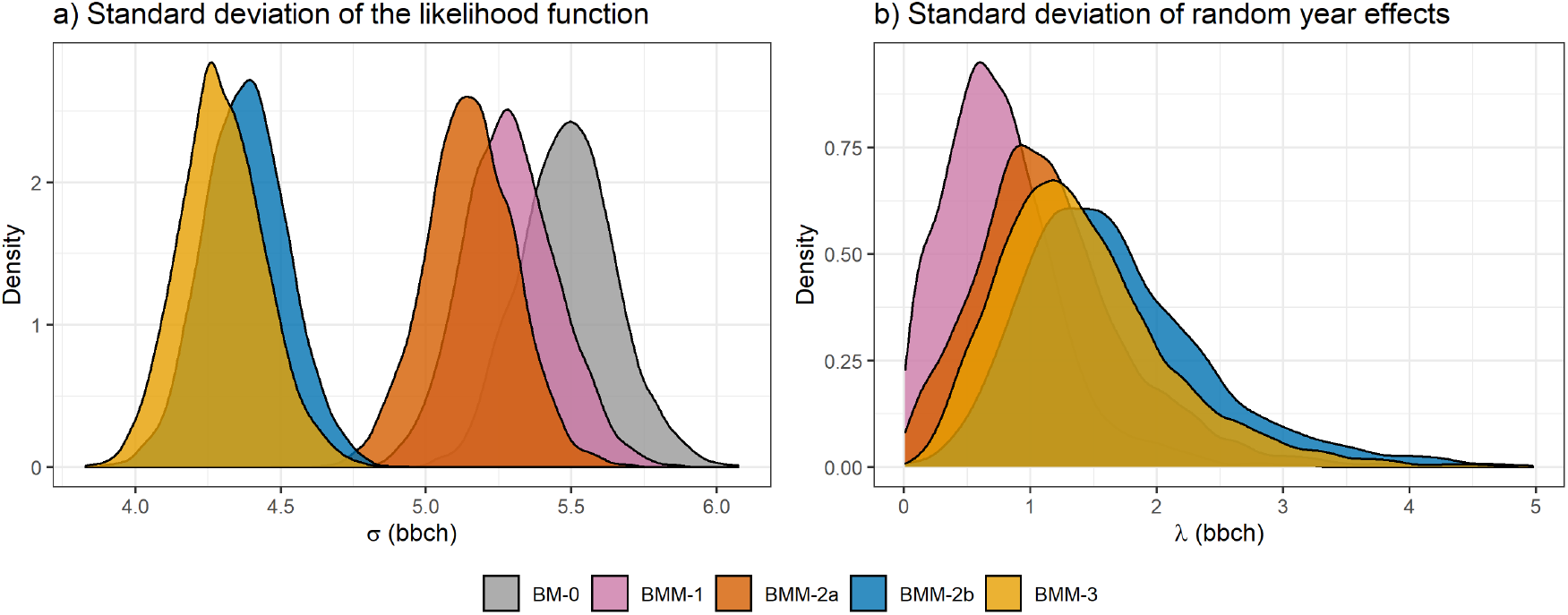
Posterior parameter distributions of (a) standard deviation of the likelihood function (σ) and (b) standard deviation of the random year effects (λ), for the Bayesian models (BM-0 to BMM-3), expressed in units of phenological development (bbch).

## 4 Discussion

The Bayesian multi-level models were able to improve calibration by partitioning some of the residual uncertainty into those arising from the hierarchical classification of cultivars and differences in ecoregions, weather conditions and year of growth. As seen in Fig.4, the unresolved residual uncertainty in simulated phenology was smaller in the full multi-level model case (BMM-3 in Fig.4b) than in the pooled case (BM-0 in Fig.4a). This was also shown by the reduction in the standard deviation of the likelihood function (σ) in Fig.11a. Moreover, the multi-level models reduced bias and RMSE (Fig.6), especially in the cases of BMM-2b and BMM-3. The differences between different cultivars and ripening groups were taken into account by the hierarchical structure of cultivars nested within ripening groups in BMM-2b and BMM-3. As a result, the uncertainty in simulated phenology (Fig. 4) originating from the process model parameters did not collapse as seen in the pooled case (BM-0), due to a collapse of the posterior parameter distribution (not shown). The over-confidence in the parameters of the pooled case can lead to poor predictions [27] because most of the variability between site-years, which can be attributed to cultivar and ripening group differences, are attributed to random noise (*σ*). The additional inclusion of the weather effects (from BMM-2b to BMM-3) further improved calibration quality (Fig. 6b). The full multi-level model BMM-3 allowed for a more representative estimate of SPASS model parameter uncertainty. This was seen from the wider ranges (blue bands) of the resultant simulated phenology in Fig.4b as compared to Fig. 4a.

Based on their between-cultivar variability, the posterior distributions of SPASS model parameters were grouped into: cultivar-specific (high variability) and ripening group-specific (low variability) parameters (Fig. 7). The parameters that exhibited low between-cultivar variability such as *tminv, tminr*, and *toptr* (Fig. D.13) define the Temperature Response Function (TRF). In general, cardinal temperatures are expected to be ripening group-specific, while parameters such as *pdd1* and *pdd2* (Fig. D.14), should be cultivar-specific because they represent traits that would be optimized in different cultivars by plant breeders [58, 59, 60]. Most model parameters exhibited this expected behaviour, with the exception of *toptv.* The posterior distribution of parameter *toptv*, which influences vegetative development, may not only represent the ripening group-specific optimum temperature but may have also compensated for a missing cultivar-specific effect. We assumed no photoperiod effect on vegetative development in the model since this effect is small for maize grown in temperate regions [61]. Nonetheless, such an effect could exist and be cultivar-dependent. This missing effect could have been compensated by *toptv* in the model. The base temperature for emergence *emt* also exhibited some between-cultivar variability. This parameter could have compensated for the effects of some cultivar-specific hard-coded parameters in the emergence equation (eq. A.1). Furthermore, the ripening group-level distributions of parameter *tminv* (Fig. 7a) showed that, as expected, early/mid-early ripening groups on average have a lower minimum temperature requirement for vegetative development than the mid-late/late ripening groups. Overall, the match between the behaviour of calibrated parameters and theoretical expectation highlights the model’s robustness if weather and eco-region effects are also accounted for through the multi-level modelling approach.

Analyses of the weather effects parameters highlight model deficits related to temperature effects during reproductive development. Firstly, the non-zero values of these parameters indicate that weather effects are influential (Fig. 8). Also, the narrow posterior distributions as compared to the prior show that the parameter values are informed by the observations. Furthermore, there was a high correlation between median weather effects and average reproductive phase temperature by weather class (Fig. 9). These weather effects indicate that the model overestimates phenological development at higher late summer temperatures and underestimates at lower temperatures. This trend reduced with the introduction of the cultivar-ripening group hierarchy. Certain cultivars may have been selected by the farmers based on their performance in the local environmental conditions such as temperature [4, 62, 58]. Introducing the ripening groups as a level in the hierarchy could account for the differences between the groups in reaching maturity. This differentiation of ripening groups becomes evident in the reproductive phase of development during the late summer months. In the absence of cultivar-ripening group hierarchy in BMM-2a, the weather effects captured these differences, as shown in Fig. 9a). This has implications in cases where ripening group and cultivar information are not available. Regional data sets, such as those from the state office of agricultural statistics, may not contain information about the cultivars grown, but information about environmental conditions are usually more readily available. In the absence of ripening group and cultivar information, accounting for weather effects will still result in some degree of improvement in model calibration quality. Although weakened in BMM-3, the weather effect-temperature trend was not completely removed (Fig. 9b). This indicates that model deficits related to temperature effects persist in the reproductive phase. Thus, accounting for weather effects in addition to the hierarchy in BMM-3 resulted in improved site-year calibration quality in many weather classes as compared to BMM-2b (Fig. 6b). The current TRF in the reproductive phase may not sufficiently capture the true development behaviour. Different TRFs for maize phenology should be evaluated using this approach with the aim to identify a better representation of the underlying processes.

Analyses of eco-region effects parameters point to model deficits related to soil moisture. Since eco-region effects were accounted for in all the four multi-level model cases, we first provide a detailed discussion of the results here. The median values of the eco-region effects exhibited a positive increase in many regions on the inclusion of weather effects (Fig. 10a), and a negative increase on the inclusion of cultivar-ripening group hierarchy (Fig. 10c) as compared to the BMM-1 (Fig. 10a). Accounting for either one of these factors (weather effects or cultivar-ripening group hierarchy) without the other possibly resulted in an overestimation of eco-region effects. The eco-region effects compensated for these missing factors. The neighbouring eco-regions 6 and 8 (northern Lowlands), and 2 (Rhine plain), as well as 3 and 4 (central Uplands) exhibited similar trends in eco-region effects, irrespective of the model case. Eco-regions 6 and 8 and eco-regions 3 and 4 can be further grouped, thus allowing for model simplification. The distinct differences between the Lowland, plains (negative eco-region effect) and the Uplands (positive eco-region effect) could be due to distinct climatological or pedological features. The northern Lowlands are characterized by moraines that have high groundwater levels. The Rhine plain also exhibits higher groundwater productivity than the central Uplands [63]. In the full model (BMM-3), phenological development was overestimated in the northern Lowlands and Rhine plain, when only the process model parameters, weather and year effects were considered. This overestimation was then corrected by the negative eco-region effects in these regions. This could indicate a possible influence of water-logging which slows phenological development and is not accounted for in the SPASS model. Liu et al., (2021) [64] found that accounting for water-logging stress on phenology and yield in the APSIM model improved model performance for barley. A similar consideration in the SPASS model equations may be required to account for this stress. This effect could also occur because high soil-water content or water logged soils might lower temperatures within the crop canopy through the cooling effect of evaporation, changes in albedo at the soil surface, enhanced soil heat capacity, and heat transport resulting in a heat transfer away from the soil surface. None of these effects are accounted for in the model. An alternate formulation of the multi-level model that separately accounts for factors like soil, climate and topography instead of eco-regions is suggested to aid further investigation. By separating the effects of ecological regions from the process model parameters we were able to gain further insights into the model and identify possible model deficits.

As is expected, the unresolved model residual error represented by the standard deviation of the likelihood function reduced with increasing model complexity. A large reduction was seen due to the inclusion of the ripening group-cultivar hierarchy. This could have been a consequence of including a large number of parameters corresponding to 4 ripening groups and 66 cultivars in models BMM-2b and BMM-3. The other nuisance parameter, the standard deviation of the random year effects, increased with increasing model complexity from BMM-1, BMM-2a to BMM-2b, followed by a slight reduction in BMM-3. Accounting for other effects resulted in a better estimation of the random year effects that were previously attributed to the unresolved residual error.

While Bayesian multi-level modelling improved calibration, we acknowledge the limitations of our approach in using limited data and excluding input uncertainty. Although the inclusion of only ecoregion, weather and year effects do account for some improvement in model performance (Fig. 5), this was not clearly evident when model performance was analysed by eco-region and weather classes (Fig. C.12). These effects may be convoluted by a possibly stronger effect of cultivar-ripening group hierarchy, through compensation when it is not explicitly taken into account. Although, more observation data would help in disentangling the different effects, they may not be available. Additionally, as noted in this study, using more data is also accompanied by high computation costs, especially for modelling cases with higher complexity. Importantly, the uncertainty in phenology observations and in the model inputs like temperature or reported sowing and harvest dates was not considered and could further confound results.

The resultant posterior distributions from the full model case (BMM-3) facilitate phenology predictions for cultivars, ripening groups and for the maize species grown in different environmental conditions in Germany. The phenological development of a current cultivar which has been used for calibration, can be predicted in a different ecological region or weather class. In this case, the posterior parameter distribution of the cultivar can be used to simulate phenology using the SPASS model, while the corresponding parameters of the ecological region and weather effects can be used to correct the model simulations for structural deficits. The random year effects and unresolved residuals are added to represent the total uncertainty in predictions. Furthermore, cultivar-level parameters within a given ripening group can be used to simulate phenology for new cultivars in that group, while all cultivar-level parameters can be used to predict phenological development of silage maize grown in Germany in future scenarios. Although multi-level models are expected to improve prediction quality [42], this may not always occur. Fer et al., (2021)[35] showed that Bayesian hierarchical modelling does not always lead to best predictions. Instead, it may result in a better representation of prediction uncertainty. This lack of improvement in prediction quality could be attributed to bias-variance trade-off, in which we introduce more explanatory parameters in the multi-level models at the cost of over-fitting. Although there is always a danger of over-fitting, we justify the complexity of the multi-level models since we employed a systematic approach to increase complexity based on system knowledge. While assessing prediction quality of the models is important, this is beyond the current scope of gaining a better understanding of the phenology model and identifying model deficits in its application to regional studies. The BMM approach can also be applied to process-based crop models, wherein these point-based models can be spatialized [65]. Additionally, gene-based models can be integrated with crop models to determine more representative genotype-specific parameters [66, 67]. Thus, BMM can also be used a diagnostic tool to guide model improvement efforts.

## 5 Conclusions

In this study, we demonstrated that Bayesian multi-level modelling (BMM) is a suitable approach to account for the hierarchical structure of cultivars nested within ripening groups of a crop species, while simultaneously providing insights into model deficits related to environmental factors. Accounting for the eco-regions, weather and year effects, and specifically the hierarchical classification of cultivars and maize ripening groups led to better calibration and representation of parameter uncertainty as compared to the commonly used pooled approach. With our approach, models that have been primarily used for field scale or cultivar-specific studies can be extended to regional scales. Although we do not explicitly correct the process model equations, we account for effects of model deficits related to environmental conditions. This approach also highlights model deficiencies which can facilitate model improvement. These findings can be used to design dedicated experiments and data-selection procedures to support the refinement of model equations. BMM could also be applied to small-scale studies to account for between-farm and within-farm variability. BMM is a valuable tool in the Bayesian tool-box that should be implemented in crop model calibration studies.

## 6 Data availability

Phenology observations used for the study are available with the German National Meteorological Service (Deutsche Wetterdienst-DWD) [46]. The ERA5-Land re-analysis gridded data set are available at the Climate Data Store (CDS), Copernicus Knowledge Base [48]. The ecological regions were obtained from the Bundesamt für Naturschutz [49].

## 7 Code availability

The R codes used for the study are available at (A link to the Zenodo repository will be provided upon acceptance).

## 8 Acknowledgements

The contribution of Michelle Viswanathan was made possible through the Integrated Hydrosystem Modelling Research Training Group, funded by the German Research Foundation (DFG, GRK 1829). The contribution of Tobias K.D. Weber was possible through the Collaborative Research Centre 1253 CAMPOS (Project 7: Stochastic Modelling Framework), funded by the German Research Foundation (DFG, Grant Agreement SFB 1253/1 2017). The authors thank Daniela Bendel for help with the DWD data. The authors acknowledge support by the state of Baden-Wuerttemberg through the HPC cluster bwUniCluster (2.0).

## Appendices

### A SPASS model equations

Phenological development in the SPASS model occurs in three main phases: emergence, vegetative phase and reproductive phase. The rate of phenological development during emergence or the emergence rate is a function of sowing-depth (Sdep) in cm and a minimum or base temperature (emt) in °C requirement only above which emergence occurs. For a particular day *d* between sowing and harvest, emergence rate RdevE_d_ (*d*^-1^) is given by

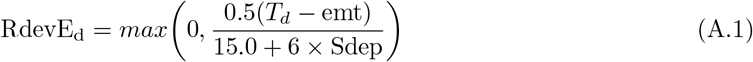

The temperature response function (TRF) defines the phenological development rate during the vegetative and reproductive phases, as a function of temperature. Development occurs only between the minimum (tmin) and maximum (tmax) temperature defined by the TRF, with maximum development occurring at the optimum temperature (topt). The TRF is given by

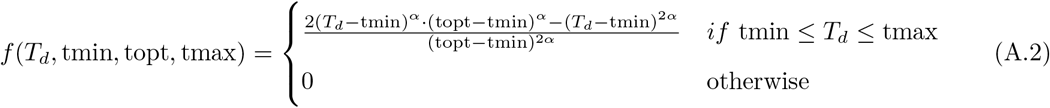

where

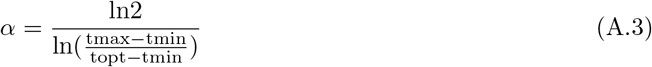

These cardinal temperatures are expressed in °C and are development phase-specific. Thus, for the vegetative phase, the development rate RdevV_d_ (*d*^-1^) is dependent on the phase-specific TRF (*fv*) (-) and the maximum development rate 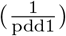 for the vegetative phase at optimum temperature (*d*^-1^). The TRF scales between 0 and 1 and acts as a reduction factor on the maximum development rate when the temperature is not at the optimum.

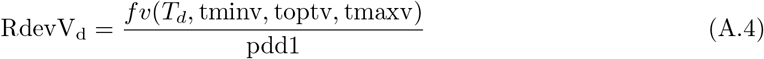

Similarly, for the reproductive phase, if 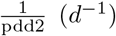 is the maximum development rate at optimum temperature and *fr* (-) is the TRF, then the development rate RdevR_d_ (*d*^-1^) is given by

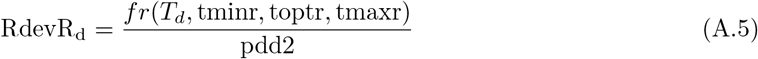

The phenological development rate *Rdev_d_* (*d*^-1^) at a given day *d* is given by

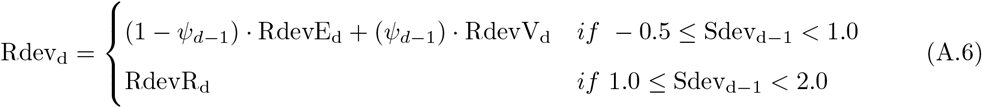

where

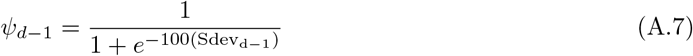

In eq.A.6 we introduced a slight modification for the original SPASS model equations [43]. We defined a sigmoid *ψ*_*d*–1_ instead of a step function for the transition between emergence rate and vegetative development rate.

The internal development stage Sdev_d_ is given by

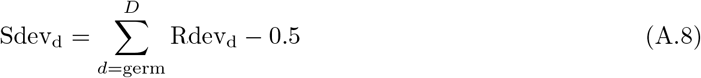

where germ is the date of germination or, in this case, date of sowing since germination is assumed to be instantaneous.

The internal development stages are converted to BBCH stages bbch_d_ by

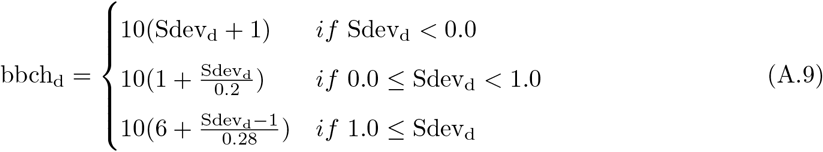

In the R and Jags implementation of the model, phenological development rate is calculated at time-step of 0.1 day. Air temperatures are first interpolated at this time-step and used as model inputs. The development rate equations A.1, A.4 and A.5 are converted from the units of (1/day) to 10/day by multiplying them by the time-step of 0.1. Phenological development is calculated from the first time-step of the sowing day to the end on the harvest day.

### B weather class clustering

The site-years available for the study, were classified into ten weather classes based on average temperatures and cumulative precipitation between April and June, and between July and September. K-means clustering was used to define the weather classes. The K-means algorithm generates clusters by minimizing within-cluster variations that are based on Euclidean distances [68]. First, the average of the mean daily temperature (*T_sy,s_*) from April to June and from July to September were calculated for each site-year (*sy*) as follows:

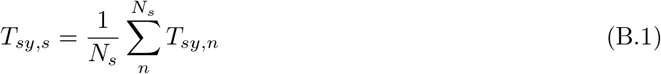

where *s* represents the season from April to June or from July to September, *N_s_* is the total number of days in that season, and *T_sy,n_* is the mean temperature (°C) on a given day *n* at site-year *sy*. Similarly, cumulative precipitation (*P_sy,s_*) was also calculated from April to June, and from July to September as follows:

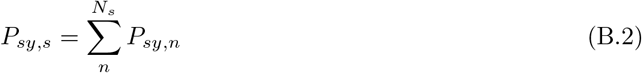

where *P_sy,n_* is the precipitation (mm) on a given day *n* at site-year *sy*.

The values for each of the four factors (*T_sy,Apr–Jun_, T_sy,Jul–Sep_, P_sy,Apr–Jun_, P_sy,Jul–Sep_*) per site-year were then normalized by scaling the values between 0 and 1.

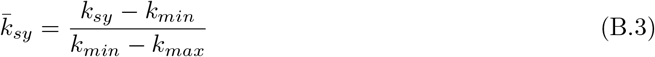

where *k_sy_* represents the factor, 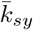 is its normalized value, and *k_min_* = *min*(*k*_1_, *k*_2_,…*k_SY_*) and *k_max_* = *max*(*k*_1_, *k*_2_,…*k_SY_*) are the minimum and maximum values of the particular factor across all the site-years (*SY* = 3, 004), respectively. The *kmeans* function from the *stats* package [69] in R was run to generate ten clusters. To ensure stability of the resultant clusters, 100 starting points were set. The maximum number of iterations was set to 1000 and 10 clusters were specified to generate 10 weather classes.

### C Model performance metrics

The model calibration performance represented by the mean RMSE and bias from the 100 site-years used for calibration (Fig. C.12), was analysed by eco-region and year for the five model cases BM-0, BMM-1, BMM-2a, BMM-2b, BMM-3. Overall, the ripening-cultivar hierarchy in BMM-2b and BMM-3 show a clear improvement in model calibration quality. The inclusion of eco-region and year effects from BMM-1 onwards, does not show a clear improvement trend across all classes.

**Figure C.12:**
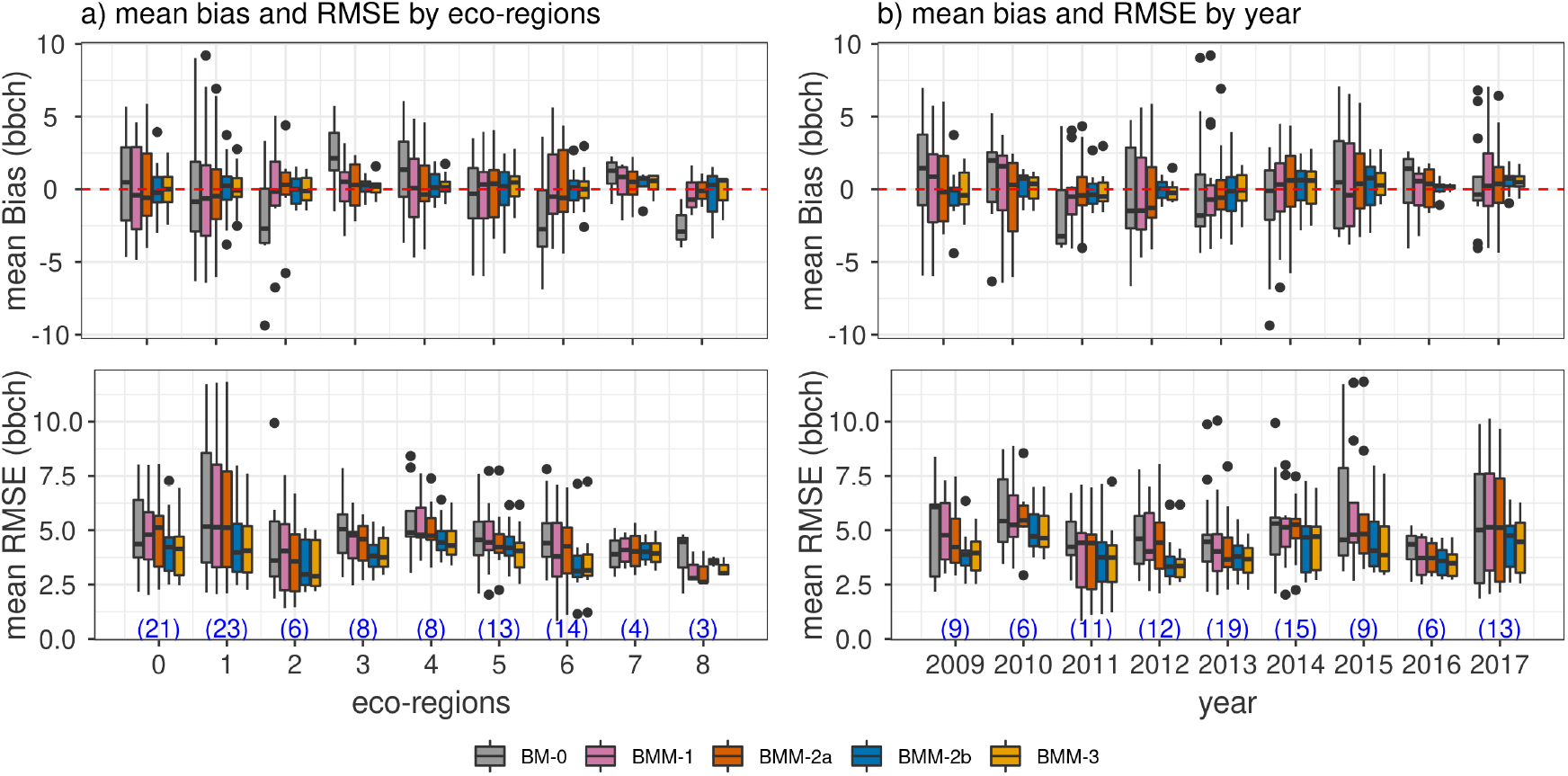
Box-plots of the mean rmse and bias by site-year within eco-regions (a), years (b) for each of the five model cases: BM-0 (pooled), BMM-1 (eco-regions and random year effects), BMM-2a (eco-regions, weather class, and random year effects), BMM-2b (cultivar-ripening group hierarchy, eco-region, and random year effects), BMM-3 (full model with cultivar-ripening group hierarchy, ecoregions, weather class, and random year effects). The numbers in blue at the bottom of each plot indicate the number of site-years and consequently the number of points in the groups defined on the x-axis that were used to obtain the box-plots. Hinges of the box-plot represent the inter-quartile range (IQR), whiskers extend from the hinges up to 1.5×IQR and values beyond this range are plotted as points.

### D SPASS model parameter distributions

Figures D.13 and D.14 show posterior distributions of parameters that exhibit low and high between-cultivar variability, respectively. In general, parameters that exhibit low between-cultivar variability, correspond to the temperature response function (TRF) in the vegetative and reproductive phases.

**Figure D.13:**
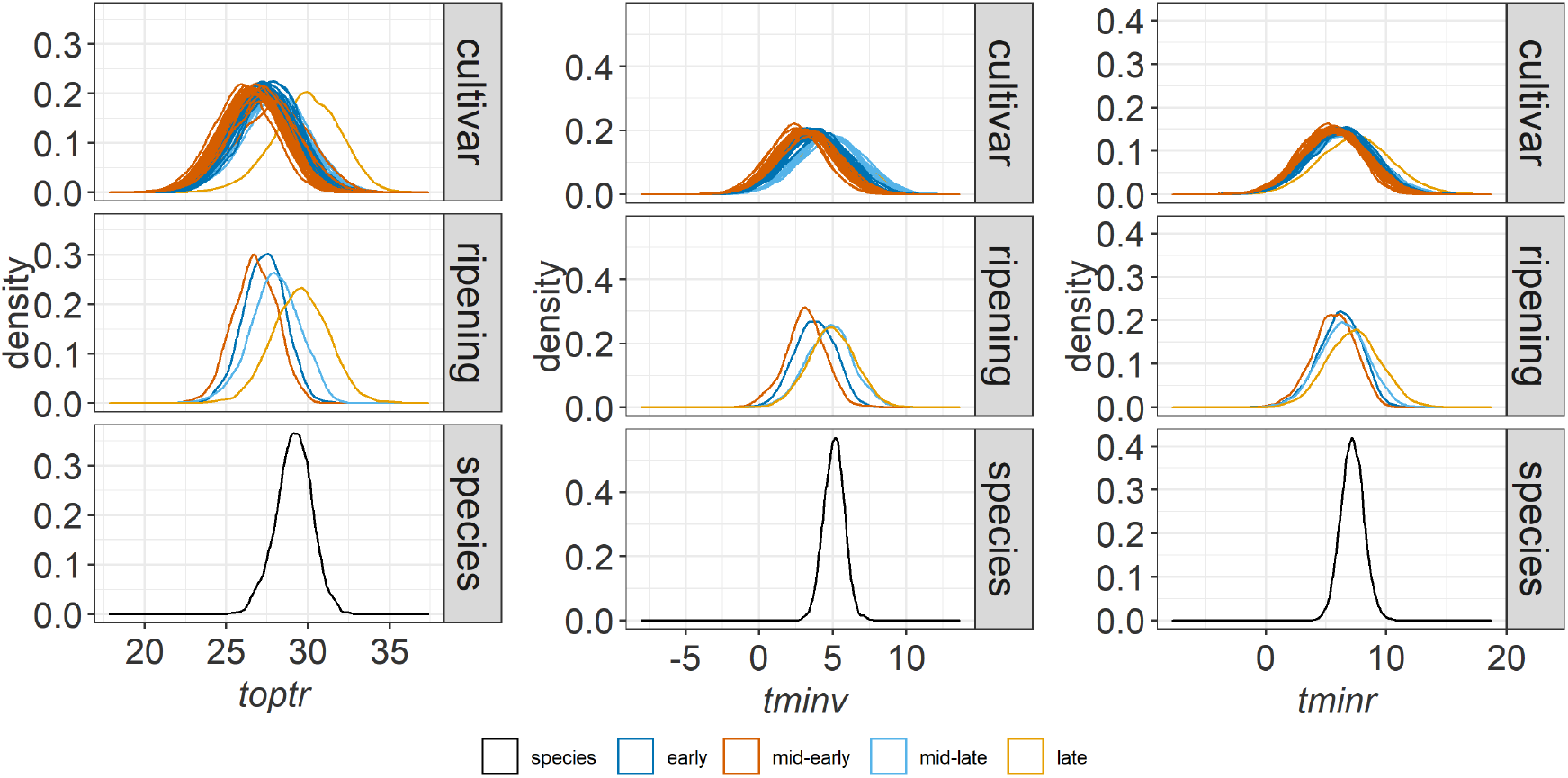
Posterior distributions of parameters that exhibit low between-cultivar variability. Parameters include: minimum and optimum temperatures for development in the reproductive phase (*tminr, toptr*) and minimum temperature for development in the vegetative phase (*tminv*) for the Bayesian full model case BMM-3. The distributions are provided for the species, ripening group and cultivar levels of the hierarchy. The cultivar distributions are coloured by their corresponding ripening groups.

**Figure D.14:**
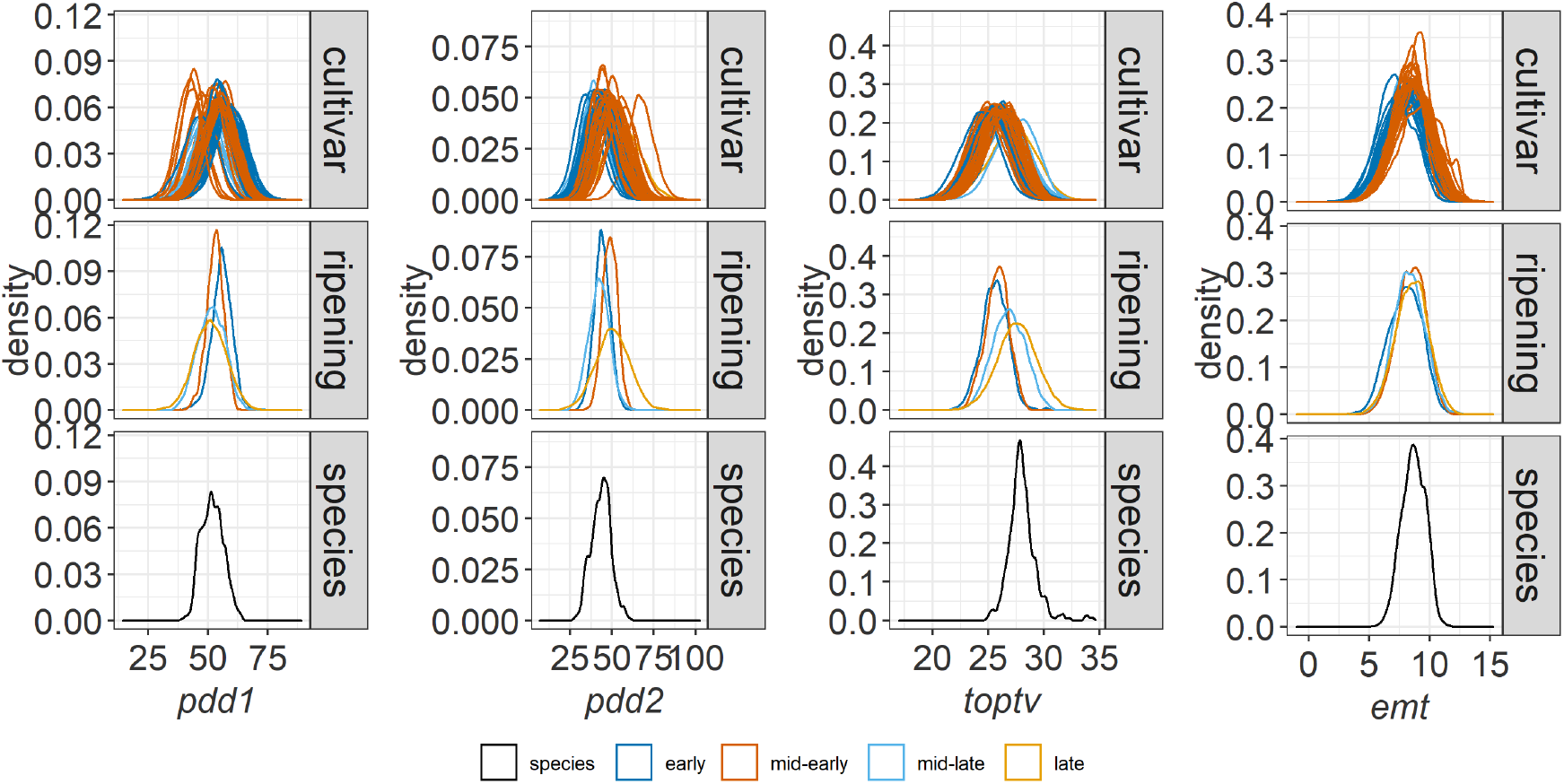
Posterior distributions of parameters that exhibit high between-cultivar variability. Parameters include: phyisiological development days for vegetative (*pdd1*) and generative (*pdd2*) phases, optimum temperature for development in the vegetative phase (*toptv*), base temperature for emergence (*emt*) for the Bayesian full model case BMM-3. The distributions are provided for the species, ripening group and cultivar levels of the hierarchy. The cultivar distributions are coloured by their corresponding ripening groups

## Notes

### Competing Interest Statement

The authors have declared no competing interest.

