## Supplementary Material for "Bayesian multi-level calibration of a process-based maize phenology model"

**S2 Dataset information** **16**

**S3 Species-level parameter distributions** **18**

**S4 Posterior parameter distributions - BMM-2b** **19**

**S5 SPASS phenology model comparison: R vs. ExpertN-5.0** **20**

### S1 MCMC posterior samples

Markov Chain Monte Carlo (MCMC) sampling of the posterior parameter distributions was performed using the R2jags [1] (for cases BM-0, BMM-1, BMM-2a and BMM-2b) and jagsUI [2] packages (for BMM-3) in R [3]. In the following section, diagnostic plots are provided for some MCMC parameter samples from the five model cases. The SPASS model parameters at the species level ($\theta_{sp}$) of the hierarchy are emt_sp, pdd1_sp, tminv_sp, toptv_sp, pdd2_sp, tminr_sp, and toptr_sp for model cases BM-2b and BMM-3, and emt, pdd1, tminv, toptv, pdd2, tminr, and toptr for model cases BM-0, BMM-1, and BMM-2a. The standard deviation of the likelihood function is sigma ($\sigma$). Parameters weath, eco, and year correspond to the weather effects ($\delta_w$), eco-region effects ($\gamma_e$), and year effects ($\tau_y$), respectively.

#### S1.1 Trace-plots

The trace-plots and density plots (coda package [4] in R [3]) for some parameters from the BMM-2a case are provided as an example. In Fig. S1 parameter sigma ($\sigma$) shows good mixing across the three chains, while parameters pdd1 and pdd2 show relatively poor mixing. The poor mixing is attributed to the parameter correlations (section S1.4). Parameters weath ($\gamma_w$) (Fig. S2), eco ($\gamma_e$) (Fig. S3), and year ($\tau_y$) (Fig. S4) show good mixing.

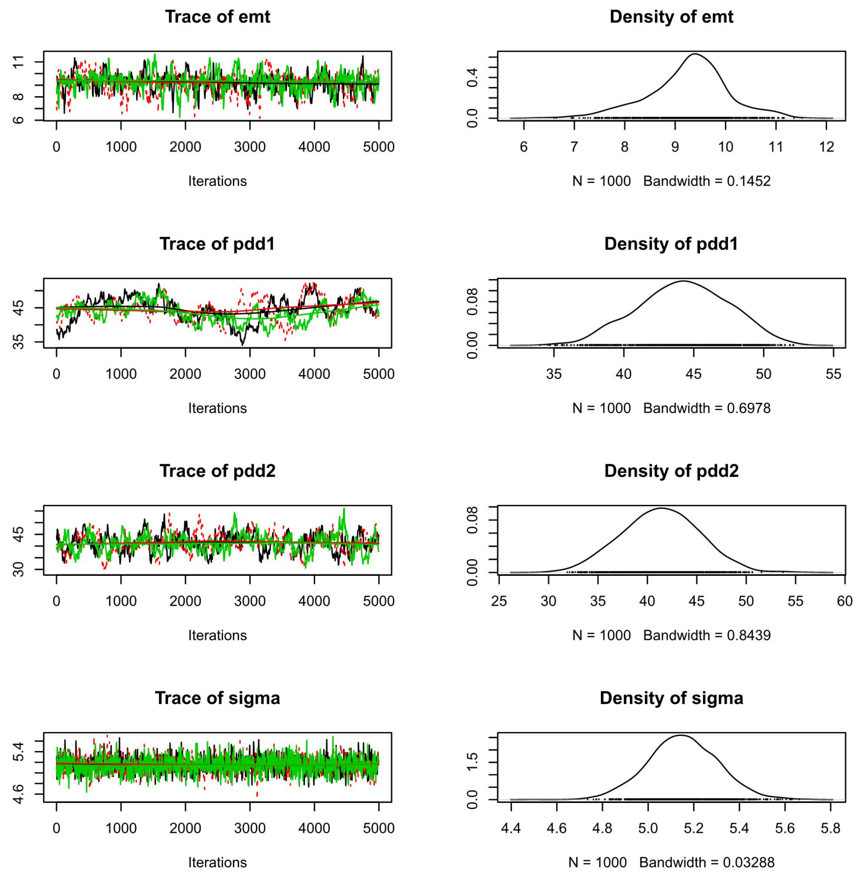

Figure S1: Trace-plots (left) and density plots (right) for some of the species level SPASS model parameters and the standard deviation of the likelihood function from the BMM-2a case.

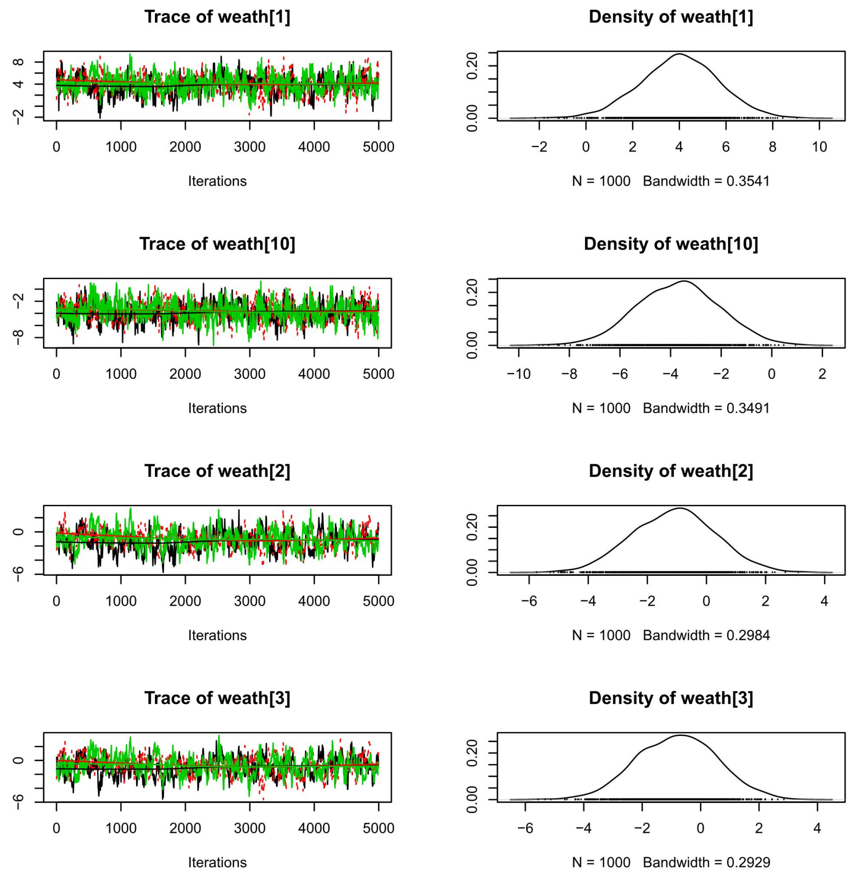

Figure S2: Trace-plots (left) and density plots (right) for weather effect parameters for some of the weather classes from the BMM-2a case.

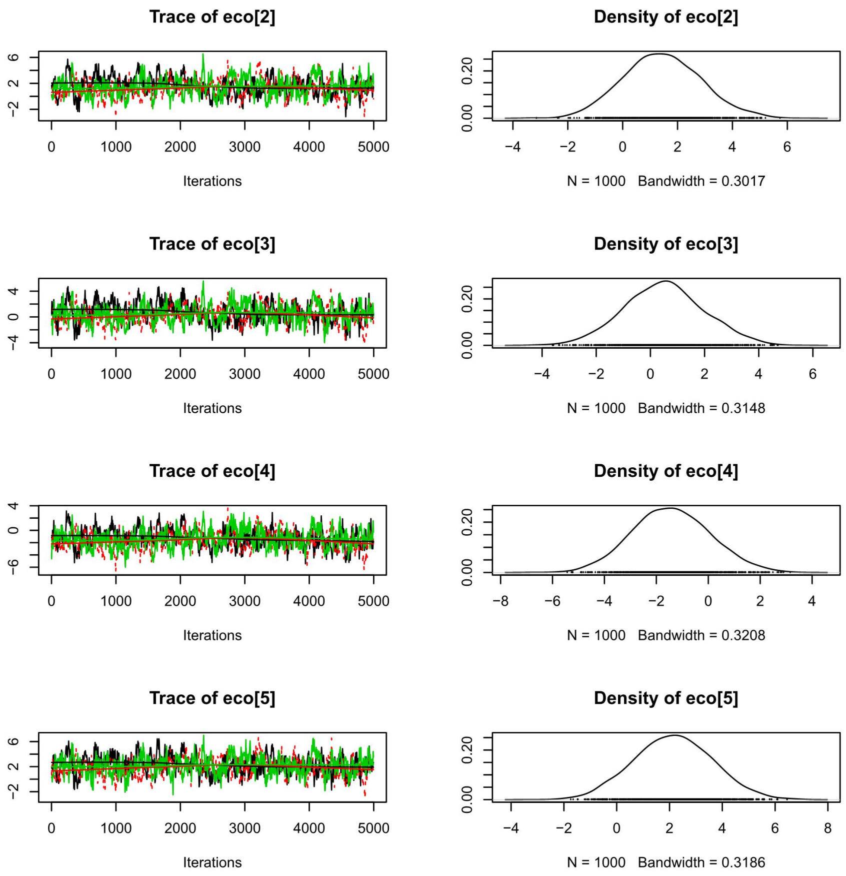

Figure S3: Trace-plots (left) and density plots (right) for the eco-region effect parameters for some of the eco-regions from the BMM-2a case.

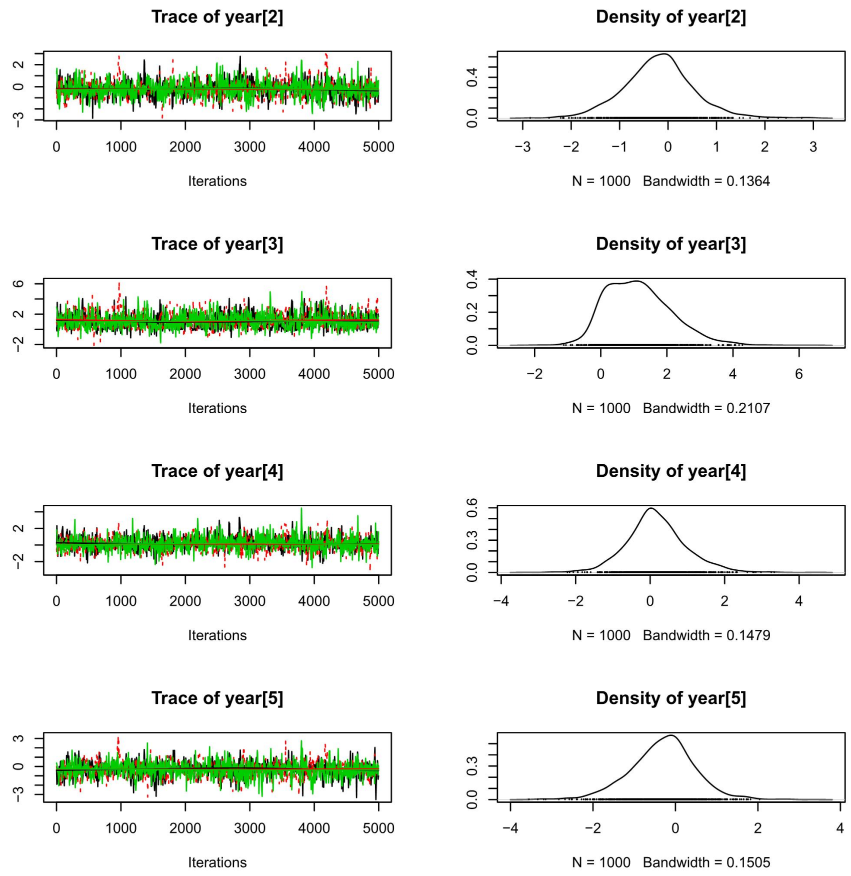

Figure S4: Trace-plots (left) and density plots (right) for the year effect parameters for some of the years from the BMM-2a case.

#### S1.2 Convergence diagnostic

The three chains for the MCMC algorithm were run until the Gelman-Rubin convergence diagnostic was $\leq 1.1$. In Fig. S5 we provide a plot of the shrink factor or the convergence diagnostic (gelman.plot in coda package [4]) for some of the parameters from the BMM-3 case. It can be seen that the parameters have converged after 3,500 iterations.

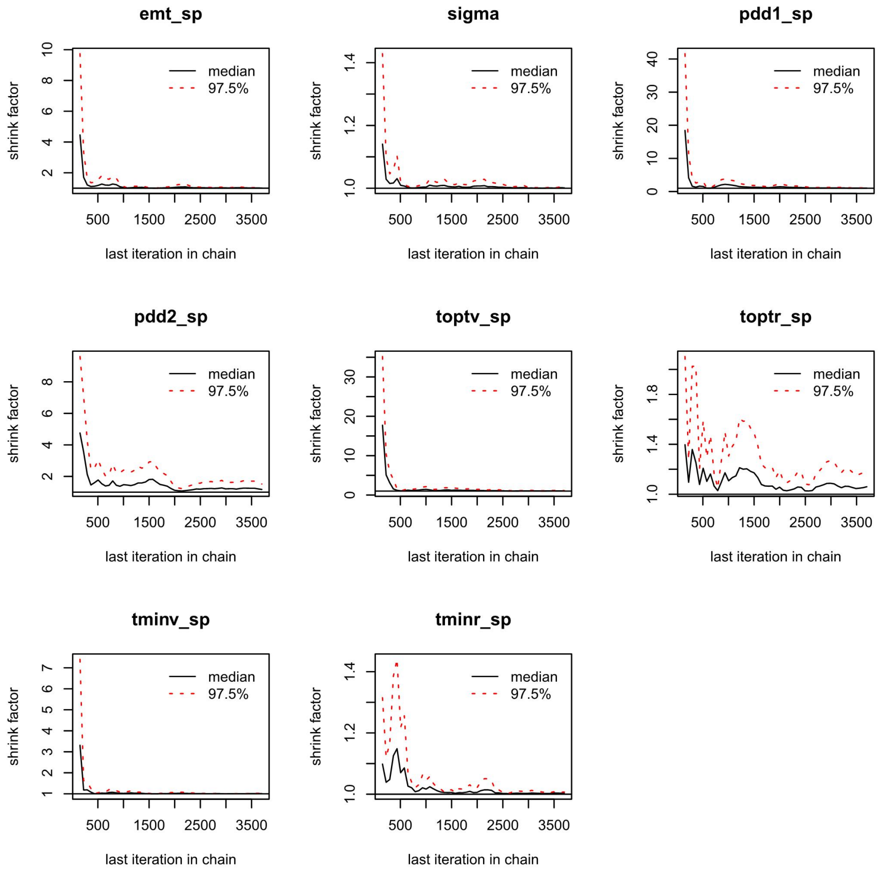

Figure S5: Evolution of Gelman-Rubin shrink factor (y-axis) for some species-level parameters ($\theta_{sp}$) in BMM-3 case with an increase in the number of iterations (x-axis)

#### S1.3 Auto-correlation plots

The auto-correlation plots (acfplot in coda package [4]) are provided for the species-level parameters ($\theta_{sp}$) and sigma ($\sigma$) in the BM-0 (Fig. S6) and BMM-3 (Fig. S7) cases. These show the auto-correlation of the parameters within the chain. The auto-correlation decreases to zero with greater lag. Note that in BM-0 the 5,000 posterior samples of were thinned to obtain a final set of 1000 samples. The samples were not thinned in BMM-3. Parameters toptv_sp ,toptr_sp, pdd1_sp, and pdd2_sp exhibit a higher auto-correlation which can be attributed to between-parameter correlations (section S1.4.

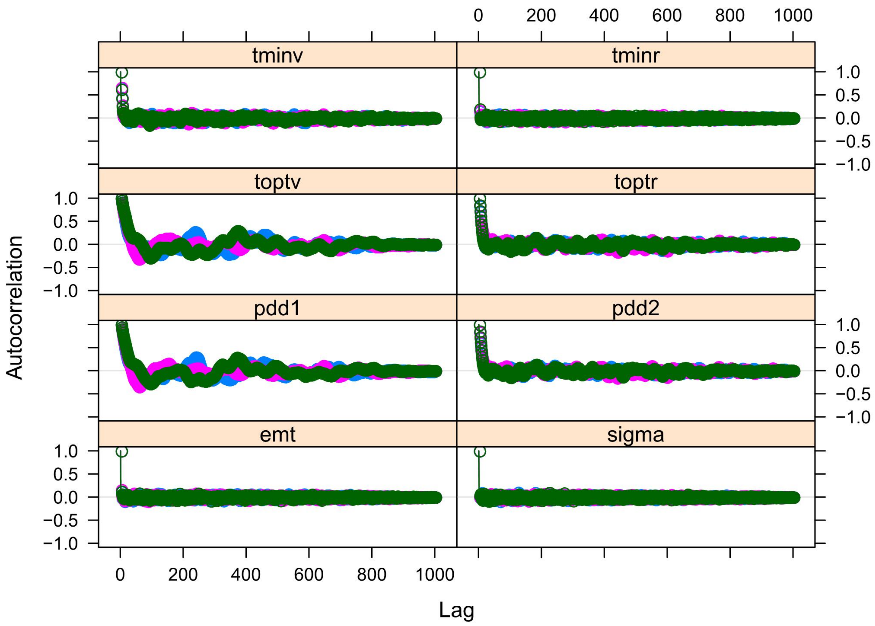

Figure S6: Auto-correlation plots for some parameters in the BM-0 case.

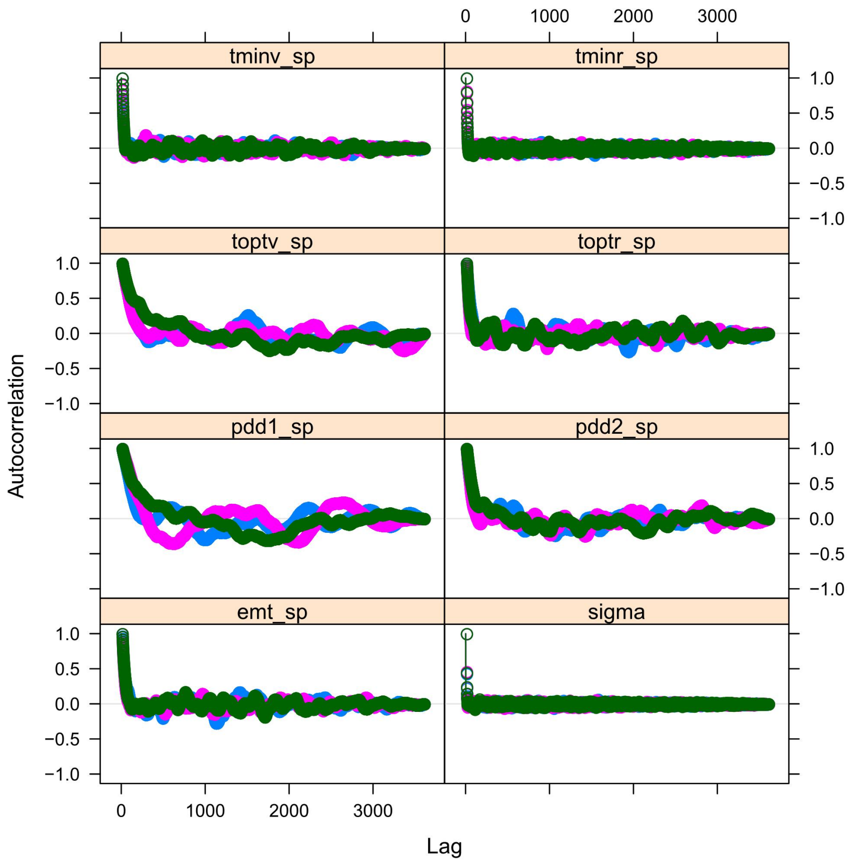

Figure S7: Auto-correlation plots for some parameters in the BMM-3 case.

#### S1.4 Correlation plots

Correlation plots (ipairs function in IDPmisc package [5]) of the species-level parameters ($\theta_{sp}$) in the hierarchy and sigma ($\sigma$) of the likelihood function for cases BM-0 and BMM-2b are provided in Fig. S8 and S9. Additionally, correlation coefficient plots (corrplot function in corrplot package [6]) of some of the parameters are provided for all the cases (Fig. S10, S11, S12, S13, S14). The colours of the ellipse indicate positive (blue) or negative correlation (red), while the colour intensity and shape of the ellipse indicates the value. A correlation coefficient of one is a diagonal line, while no correlation is represented by a white circle. There is strong negative correlation between parameters pdd1 and toptv as well as between pdd2 and toptr in BM-0 (Fig. S8), but this is not seen in BMM-2b(Fig. S9). The correlation coefficient plots show that in BM-0 (Fig. S10), BMM-1 (Fig. S11), and BMM-2a (Fig. S12), the correlation exists but is not seen in BMM-2b (Fig. S13) and BMM-3 (Fig. S14) where the ripening and cultivar hierarchy is introduced. In these two cases, these correlations are seen in the ripening- and cultivar-specific parameters (not shown). The eco-regions effect parameters (eco) are positively correlated with each other and also to the base temperature for emergence (emt) (Fig. S11). The weather effect parameters (weath) in BMM-2a (Fig. S12) are also positively correlated with each other and negatively correlated with the eco-region effect parameters (eco). This correlation between eco-region and weather effect parameters could be because there is a very likely overlap between the weather classes and eco-region class as the eco-regions are also based on climatological characteristics.

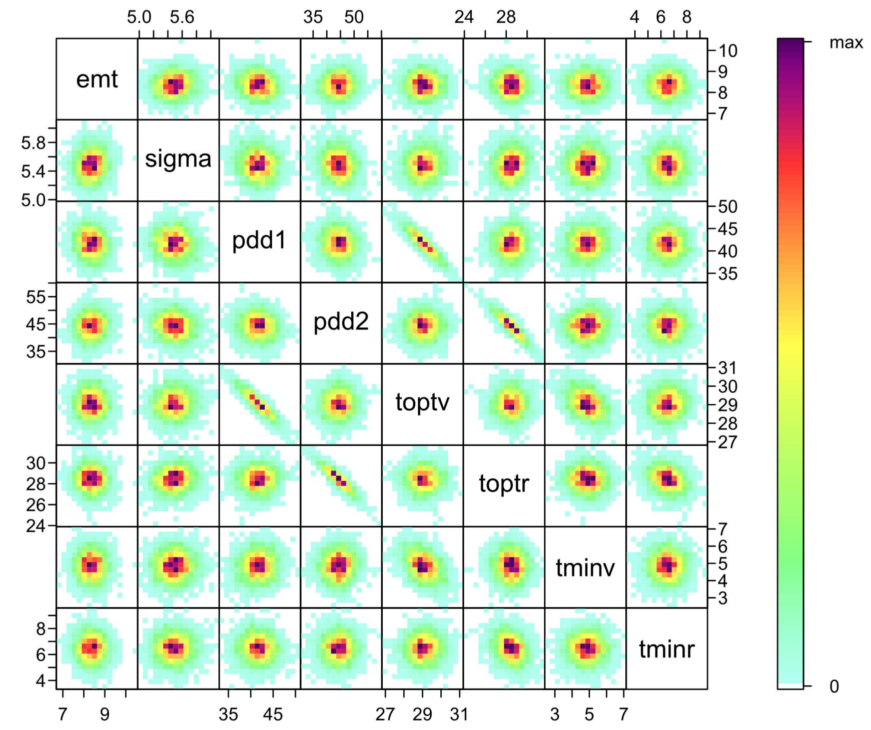

Figure S8: Cross-plot of the posterior samples of the some estimated parameters in the BM-0 case. Red represents high density and blue low density.

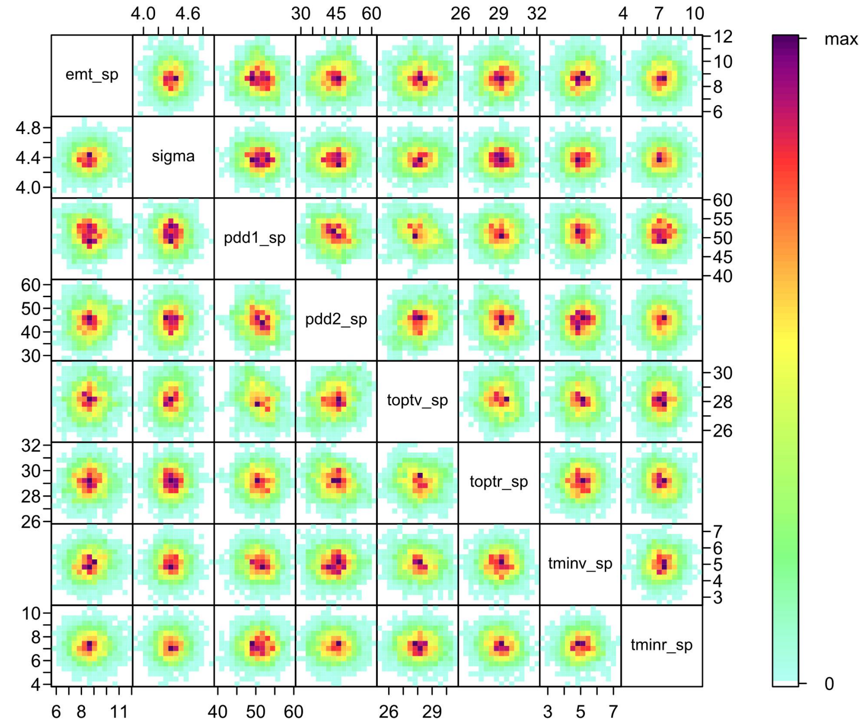

Figure S9: Cross-plot of the posterior samples of the some estimated parameters in the BMM-2b case. Red represents high density and blue low density.

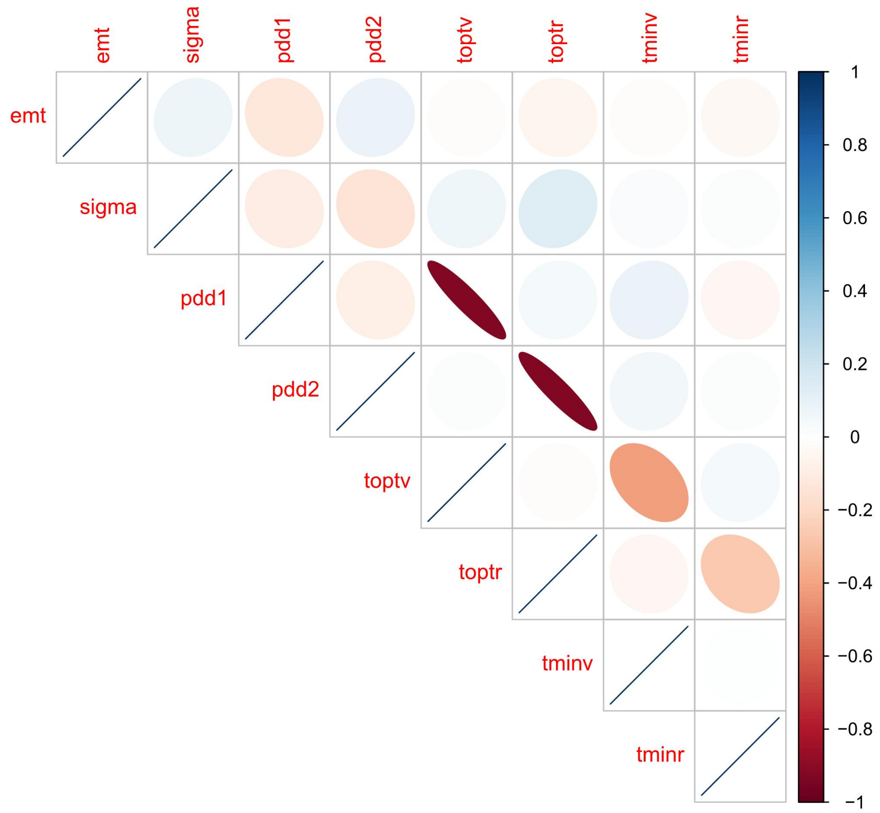

Figure S10: A plot of the correlation coefficients between some of the parameters in the BM-0 case. The colours of the ellipse indicate positive (blue) or negative correlation (red), while the colour intensity and shape of the ellipse indicates the value. A correlation coefficient of one is a diagonal line, while no correlation is represented by a white circle.

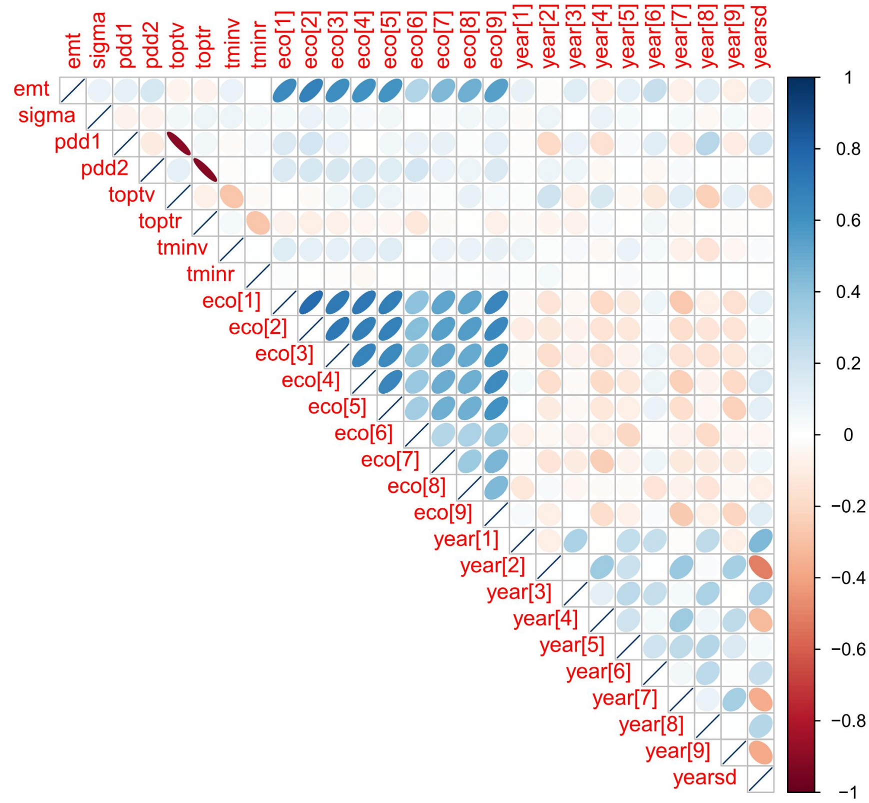

Figure S11: A plot of the correlation coefficients between some of the parameters in the BMM-1 case. The colours of the ellipse indicate positive (blue) or negative correlation (red), while the colour intensity and shape of the ellipse indicates the value. A correlation coefficient of one is a diagonal line, while no correlation is represented by a white circle.

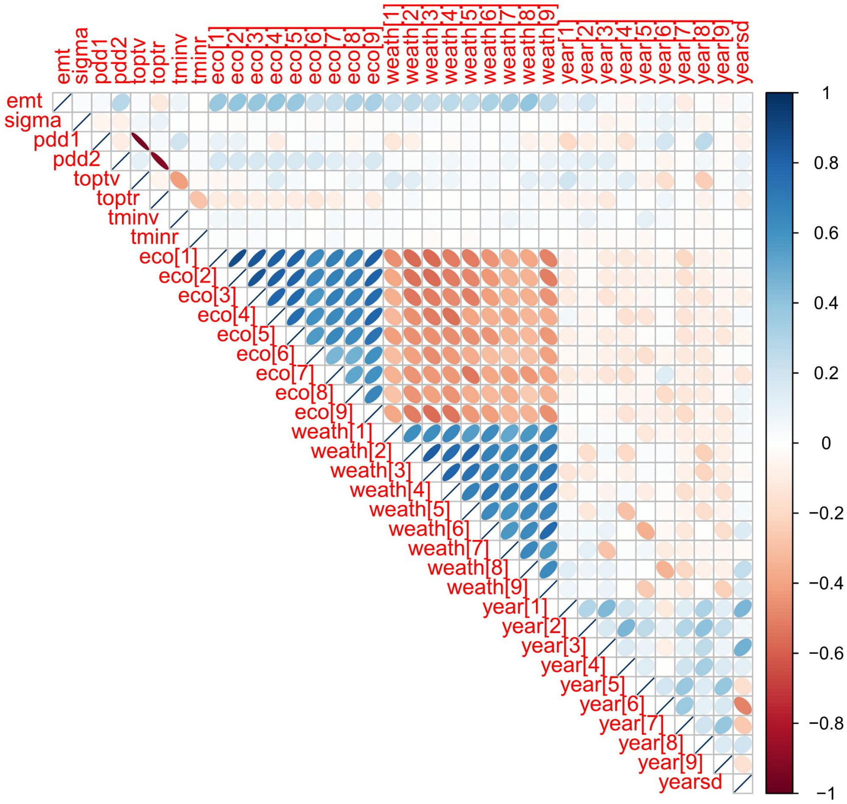

Figure S12: A plot of the correlation coefficients between some of the parameters in the BMM-2a case. The colours of the ellipse indicate positive (blue) or negative correlation (red), while the colour intensity and shape of the ellipse indicates the value. A correlation coefficient of one is a diagonal line, while no correlation is represented by a white circle.

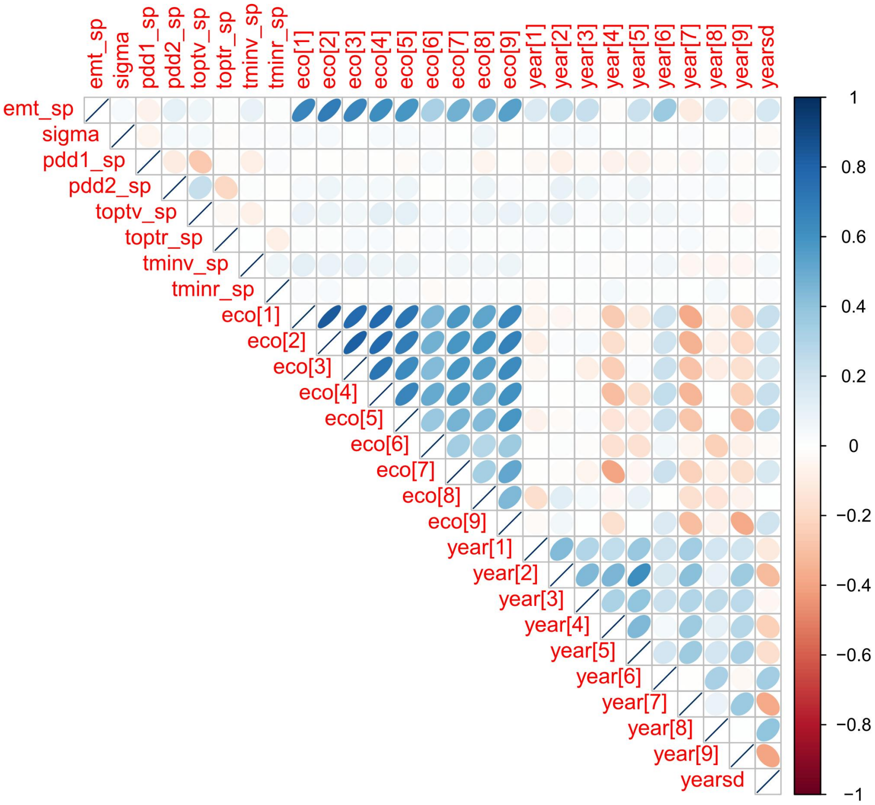

Figure S13: A plot of the correlation coefficients between some of the parameters in the BMM-2b case. The colours of the ellipse indicate positive (blue) or negative correlation (red), while the colour intensity and shape of the ellipse indicates the value. A correlation coefficient of one is a diagonal line, while no correlation is represented by a white circle.

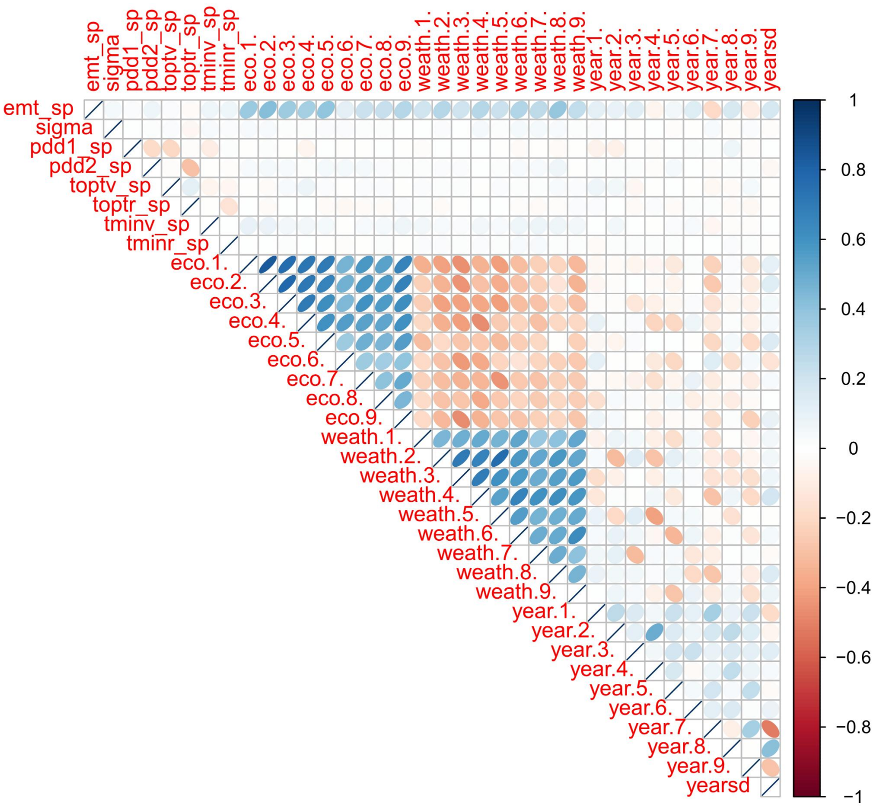

Figure S14: A plot of the correlation coefficients between some of the parameters in the BMM-3 case. The colours of the ellipse indicate positive (blue) or negative correlation (red), while the colour intensity and shape of the ellipse indicates the value. A correlation coefficient of one is a diagonal line, while no correlation is represented by a white circle.

### S2 Dataset information

Figure S15 shows (plotly package [7]) the names of the different cultivars within the four ripening groups that were used for calibration. The circle represents 100 site-years used for calibration. The late ripening group (L) contains only one site-year belonging to cultivar MAS 40F.

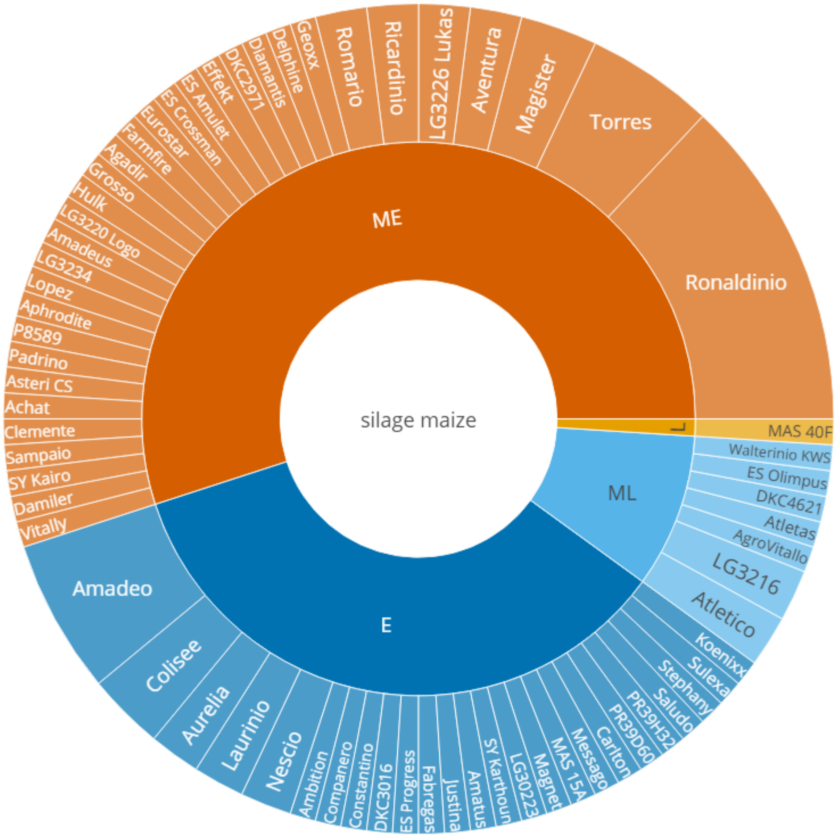

Figure S15: Cultivars and ripening groups of the 100 site-years that were used for calibration. The colours represent the ripening groups. E: early, ME: mid-early, ML: mid-late, and L: late. The circle represents 100 site-years. The cultivar MAS 40F in the late ripening group has only one site-year.

Figure S16 shows the average daily temperature and average cumulative precipitation per eco-region for April-June and July-September, based on all 3,004 site-years. The months of April-June and July-September correspond to time when vegetative and reproductive phenological development usually occurs for silage maize grown in Germany. These averages are based on 689 locations across Germany and nine years (2009-2017). Eco-region 8 (eastern part of the northern plains) has the highest temperature during April-June and July-September. In general, the southern regions received more precipitation in April-June than the northern regions. Eco-region 0 (region to the north of the Alps) received the most precipitation while eco-region 8 received the least in both periods.

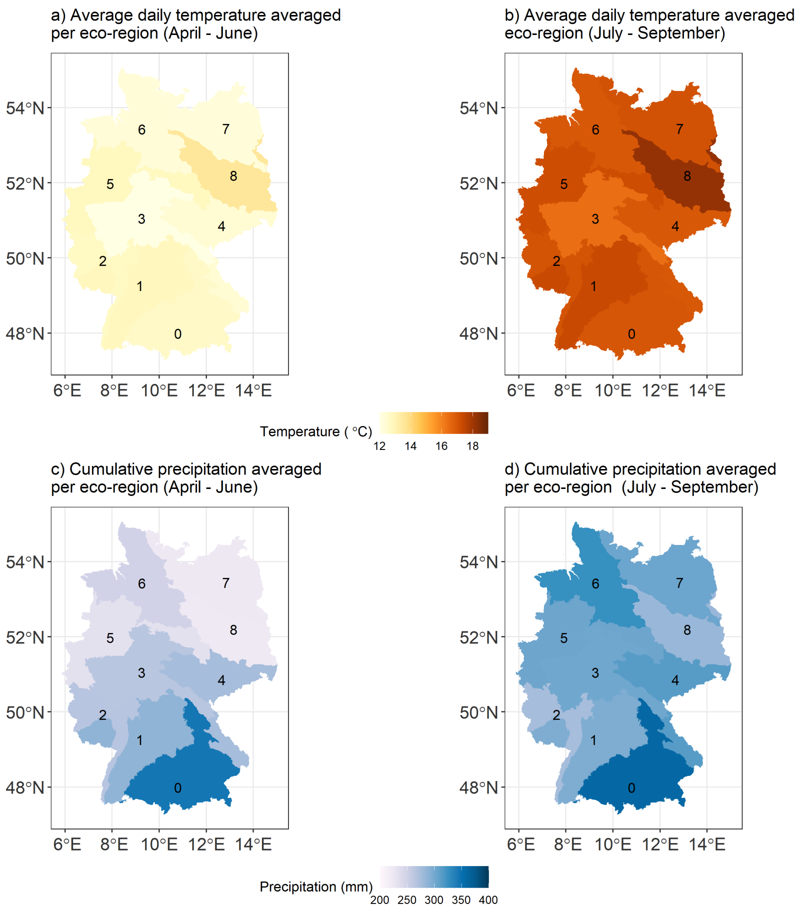

Figure S16: Average daily temperature (a, b) and average cumulative precipitation (c, d) per eco-region for April-June and July-September, based on all 3004 site-years. The numbers indicate the ecological regions based on the classification provided by the Bundesamt für Naturschutz [8].

### S3 Species-level parameter distributions

Figure S17 shows the prior and posterior distributions of species-level SPASS model parameters ($\theta_{sp}$) and the standard deviation of the likelihood function $\sigma$ from the five model cases. The marginal posterior distributions for parameters pdd1, pdd2, toptv, emt are wider when the cultivar-ripening group hierarchy is accounted for in BMM-2b and BMM-3. The marginal posterior distributions for parameters in the pooled case BM-0 are generally narrower than the other cases.

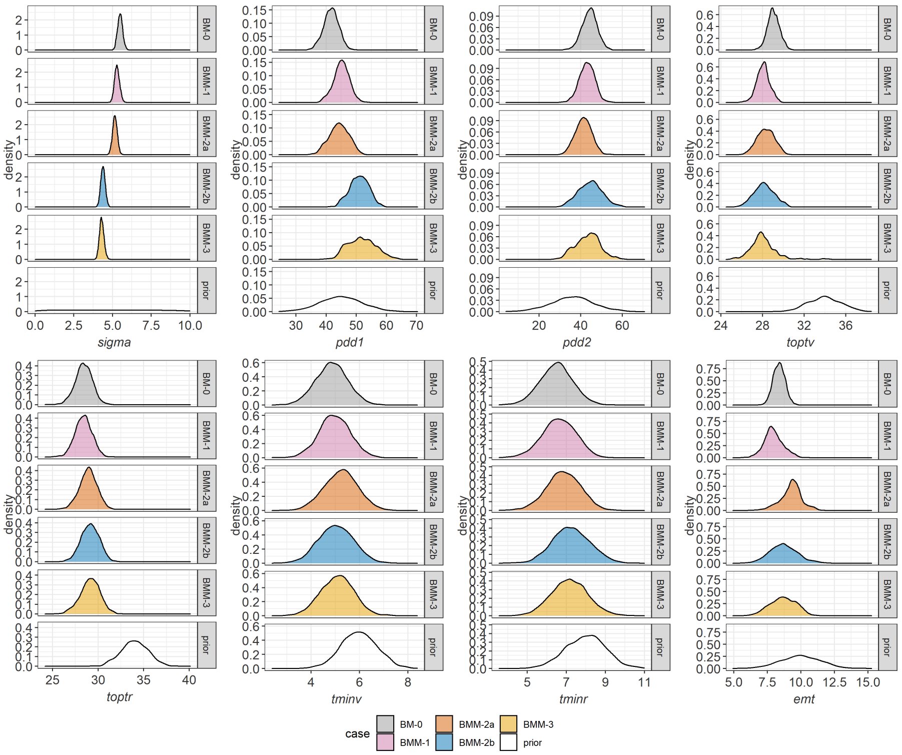

Figure S17: Prior and posterior distribution of the species-level SPASS model parameters ($\theta_{sp}$) and standard deviation of the likelihood function $\sigma$ from the five model cases BM-0, BMM-1, BMM-2a, BMM-2b and BMM-3.

### S4 Posterior parameter distributions - BMM-2b

Figures S18 and S19 show posterior distribution of parameters for BMM-2b that exhibit low and high between-cultivar variability, respectively. In general, parameters that exhibit low between-cultivar variability, correspond to the temperature response function (TRF) in the vegetative and reproductive phases, with the exception of the optimum temperature for vegetative development (toptv). The pattern is similar to the parameter distributions of in the full model case BMM-3 in the main text.

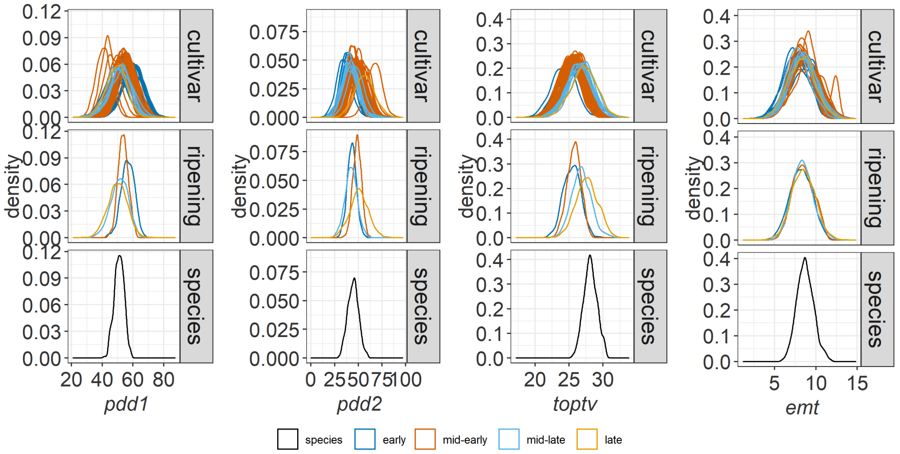

Figure S18: Posterior distribution of parameters that exhibit high between-cultivar variability. Parameters include: physiological development days for vegetative (pdd1) and generative (pdd2) phases,optimum temperature for development in the vegetative phase (toptv), base temperature for emergence (emt) for the case BMM-2b. The distributions are provided for the species ($\theta_{sp}$), ripening group ($\theta_{sp,r}$) and cultivar ($\theta_{sp,r,c}$) levels of the hierarchy. The cultivar distributions are coloured by their corresponding ripening groups.

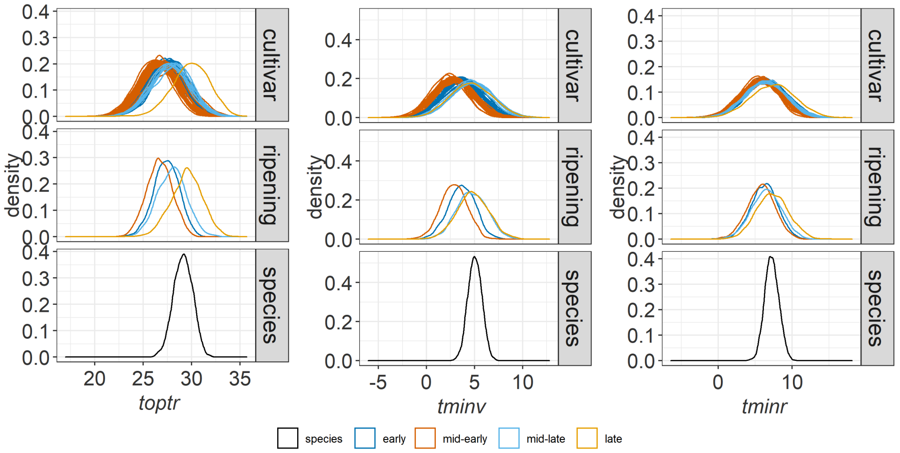

Figure S19: Posterior distribution of parameters that exhibit low between-cultivar variability. Parameters include: minimum and optimum temperatures for development in the reproductive phase (tminr, toptr) and minimum temperature for development in the vegetative phase (tminv) for the case BMM-2b. The distributions are provided for the species ($\theta_{sp}$), ripening group ($\theta_{sp,r}$) and cultivar ($\theta_{sp,r,c}$) levels of the hierarchy. The cultivar distributions are coloured by their corresponding ripening groups.

#### S5 SPASS phenology model comparison: R vs. ExpertN-5.0

Figure S20 shows a comparison between the SPASS phenology model implemented in R for the current study and the implementation in ExpertN-5 (XN5) [9] and the for a given parameter vector (pdd1=47, pdd2=43, tminv=6, toptv=28, tmaxv=44, tminr=6, toptr=28, tmaxr=44, emt=10). This difference is attributed to the sigmoidal function instead of the step function that is used for the transition between emergence and vegetative development in the current model. The differences between the two implementations are higher when phenological development is low. Overall, the differences are small.

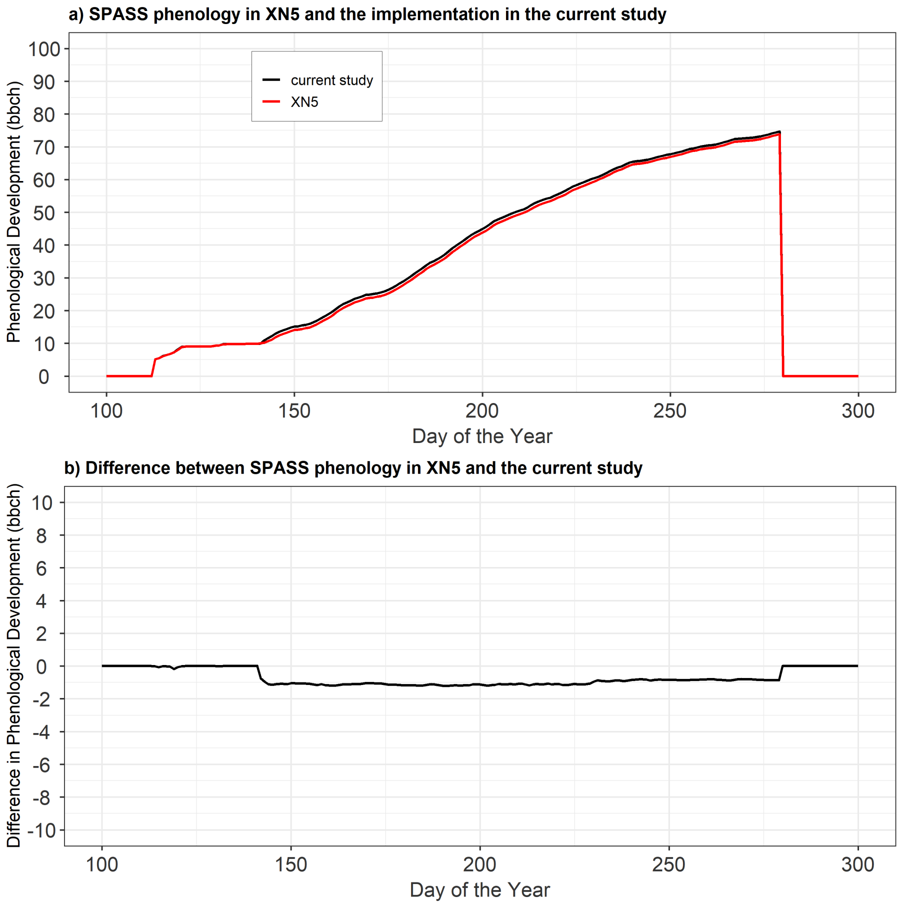

Figure S20: (a) Simulated phenological development using the SPASS model from XN5 (red) and the implementation in R (black) for the current study, for a given parameter vector. (b) Difference between the XN5 and R implementation of phenology. The current R implementation results in higher values of phenological development than the XN5 implementation. This difference is caused due to different equations being used to model the transition between emergence and vegetative development.

[2] Ken Kellner. *jagsUI: A Wrapper Around 'rjags' to Streamline 'JAGS' Analyses*, 2021. R package version 1.5.2.

[3] R Core Team. *R: A Language and Environment for Statistical Computing.* R Foundation for Statistical Computing, Vienna, Austria, 2021.

[4] Martyn Plummer, Nicky Best, Kate Cowles, and Karen Vines. Coda: Convergence diagnosis and output analysis for mcmc. *R News*, 6(1):7–11, 2006.

[5] Rene Locher. *IDPmisc: 'Utilities of Institute of Data Analyses and Process Design (www.zhaw.ch/idp)'*, 2020. R package version 1.1.20.

[6] Taiyun Wei and Viliam Simko. *R package "corrplot": Visualization of a Correlation Matrix*, 2017. (Version 0.84).

[7] Carson Sievert. *Interactive Web-Based Data Visualization with R, plotly, and shiny.* Chapman and Hall/CRC, 2020.

[8] Naturräumliche gliederung deutschlands (nach meynen/schmithüsen)., 2017.

[9] C. Klein, F. Heinlein, S. Duan, X.and Gayler, and E. Priesack. Expert-n version 5.12, 2019.
